# Molecular insights into the effect of hexanediol on FUS phase separation

**DOI:** 10.1101/2022.05.05.490812

**Authors:** Tongyin Zheng, Noah Wake, Shuo-Lin Weng, Theodora Myrto Perdikari, Anastasia C. Murthy, Jeetain Mittal, Nicolas L. Fawzi

**Author notes:** These authors contributed equally (Tongyin Zheng, Noah Wake, Shuo-Lin Weng, Theodora Myrto Perdikari).

## Abstract

1,6-hexanediol disrupts many phase-separated condensates in cells and in test tubes. In this study, we use a combination of microscopy, nuclear magnetic resonance (NMR) spectroscopy, molecular simulation, and biochemical assays to probe how alkanediols suppress phase separation and why certain isomers are more effective. Alkanediols of different lengths and configurations are all capable of disrupting phase separation of the RNA-binding protein Fused in Sarcoma (FUS), though potency varies depending on both geometry and hydrophobicity, which we measure directly. Alkanediols induce a shared pattern of changes to the protein chemical environment though to differing extents. Consistent with the view that alkanediols disrupt phase separation driven by hydrophobic groups, they decrease the thermal stability of a model globular protein. Conversely, 1,6-hexanediol does not disrupt charge-mediated phase separation, such as FUS RGG-RNA and poly-lysine/poly-aspartic acid condensates. All-atom simulations show that hydroxyl groups in alkanediols mediate interaction with protein backbone and polar amino acid side chains, while the aliphatic chain allows contact with hydrophobic and aromatic residues, providing a molecular picture of how amphiphilic interactions disrupt FUS phase separation.

## Introduction

Cellular biochemistry is spatiotemporally tuned by intracellular compartments known as biomolecular condensates. Despite lacking a phospholipid bilayer, condensates concentrate specific but heterogeneous biomolecules through a process called phase separation (Banani *et al*, 2017; Shin & Brangwynne, 2017; Yang *et al*, 2020). The formation, dissolution, and localization of membraneless puncta are controlled by a plethora of factors, such as the presence of RNA species (Roden & Gladfelter, 2021; Shin & Brangwynne, 2017) and post-translational modifications within intrinsically disordered regions (IDRs) (Martin & Mittag, 2018; Snead & Gladfelter, 2019). Important studies on the nuclear pore complex (NPC), a multiprotein structure embedded in the nuclear envelope that serves as a permeability barrier potentially by phase separation of phenylalanine/glycine repeat domains, pioneered the use of alkanethiols to disrupt disordered domain interactions and interfere with NPC functions (Ribbeck & Gorlich, 2002). Furthermore, alkanediols were also shown to cause the dissociation of nuclear pore complex components (Shulga & Goldfarb, 2003). More recently, cytoplasmic and nucleoplasmic condensates such as stress granules (SGs) (Wolozin & Ivanov, 2019) have also been shown to be sensitive to dissolution by alkanediols (Kroschwald *et al*, 2015; Wheeler *et al*, 2016). Furthermore, alkanediols have been employed to assess the reversibility of liquid nuclear and cytoplasmic puncta (Kroschwald *et al*., 2015). For example, 1,6-hexanediol, 1,2-cyclohexanediol and 1,5-pentanediol are capable of dissolving nucleopores (Jaggi *et al*, 2003; Ribbeck & Gorlich, 2002), while 1,6-hexanediol also disrupts cytoplasmic granules (Tulpule *et al*, 2021), potentially by disrupting weak hydrophobic interactions. In yeast, when 1,6-hexanediol was used in tandem with compounds that increase cell permeability like digitonin, P granules dissolved but stress granules with a more solid-like character remained intact (Kroschwald *et al*, 2017). In mammalian cells, 1,6-hexanediol has been used to probe the liquidity of condensates including stress granules, though prolonged exposure is cytotoxic and generated abnormal cell morphologies that complicate these analyses (Wheeler *et al*., 2016). Although these studies provide compelling evidence that hexanediols and similar compounds can be used to assess the physical properties of *in vitro* reconstituted droplets and cellular condensates, the details of how these compounds perturb the molecular interactions underlying liquid-like assembly remain unclear.

FUS (Fused in Sarcoma) is an RNA-binding protein whose phase separation has been extensively characterized. FUS has been in the biophysical spotlight due to its disease-causing mutations (Naumann *et al*, 2018; Patel *et al*, 2015) and its interactions with nucleic acids (Daigle *et al*, 2013; Loughlin *et al*, 2019; Sama *et al*, 2014), poly(ADP ribose) (Altmeyer *et al*, 2015; Rhine *et al*, 2022) and transcription factors (Owen *et al*, 2021). FUS phase separation is also modulated by phosphorylation (Monahan *et al*, 2017), arginine methylation (Hofweber *et al*, 2018) and N-terminal acetylation (Bock *et al*, 2021). Under physiological conditions, FUS can shuttle between the nucleus and the cytoplasm and form interactions with other proteins through its serine-glutamine-tyrosine-glycine (SQYG) rich N-terminal disordered region, arginine-glycine (RGG) motifs (Chong *et al*, 2018) and globular RNA binding domains (Deng *et al*, 2014). The aberrant function of membraneless puncta containing FUS has been associated with pathology, particularly in neurodegenerative disease and cancer (Alberti & Dormann, 2019; Boija *et al*, 2021; Ryan & Fawzi, 2019; Trnka *et al*, 2021). This has led to an emerging area of research focused on developing potential therapeutics targeting protein phase separation (Babinchak *et al*, 2020; Klein *et al*, 2020; Mitrea *et al*, 2022; Schmidt *et al*, 2022). Although the molecular interactions holding together the condensed phase of FUS, such as hydrophobic, sp^2^/π contacts, and hydrogen bonds have been studied in detail (Murthy *et al*, 2019; Murthy *et al*, 2021; Wake *et al*, 2024; Zheng *et al*, 2020), little is known at the atomic level about how non-covalent interactions with alkanediols perturb the phase behavior of FUS. Some have suggested that 1,6-hexanediol acts as “mini-detergent” that disrupts hydrophobic contacts (Hedtfeld *et al*, 2024; Ribbeck & Gorlich, 2002) while others have hypothesized that particular alkanediol-protein interaction geometries explain observations that some alkanediol isomers are more potent at inhibiting phase separation than others (Gu *et al*, 2023). To fill this gap, we employ microscopy, solution-state NMR, molecular dynamics (MD) simulations, and biochemical assays to unravel the molecular-level details of the impact of alkanediols on the phase separation, structure, and motions of FUS. In particular, we seek to probe the hypothesis that alkanediols serve as hydrophobic disruptors of phase separation and clarify why some alkanediols are more potent than others.

## Results

### Effect of alkanediols on FUS LC phase separation

Alkanediols such as 1,6-hexanediol (1,6-HD), 2,5-hexanediol (2,5-HD), 1,4-butanediol (1,4-BD) and 1,5-pentanediol (1,5-PD) have been extensively used as FUS hydrogel melting (Lin *et al*, 2016) and phase separation prevention agents (Berkeley *et al*, 2021; Li *et al*, 2021; Liu *et al*, 2021). We sought to determine how these alkanediols as well as 1,2-hexanediol (1,2-HD) and 1,2-cyclohexanediol (1,2-CHD) alter the capacity of purified protein to form liquid droplets *in vitro* using the isolated low-complexity (LC) domain (residues 1-163) of FUS as a model. First, we tested the impact of 1,6-HD on FUS LC phase separation (0% to 5%) using differential interference contrast (DIC) (**Figure 1A**) and fluorescence microscopy (**Figure S1A**). FUS LC droplet area linearly decreases between 0% and 5% concentrations of 1,6-HD (**Figure 1B**). To complement these microscopy assays, we measure the protein saturation concentration (*C*_sat_) by centrifuging phase separated samples to pellet FUS LC droplets in the presence of different concentrations of 1,6-HD (**Figure 1C**) and then quantified the protein remaining in the supernatant by UV absorbance (**Figure S1B**), which also show a linear dependence on 1,6-HD concentration. However, by 5% 1,6-HD, no FUS LC droplets were present. Therefore, to quantitatively assess the effects of 1,6-HD at higher concentrations up to the 10% 1,6-HD concentration sometimes used in cellular studies, we used the longer FUS LC-RGG1 segment, which has a decreased *C*_sat_ (i.e. an increased ability to undergo phase separation) compared to FUS LC (Wake *et al*., 2024). Similarly, we observed that FUS LC-RGG1 *C*_sat_ linearly decreases across the 0% to 10% 1,6-HD concentration range (**Figure S1C**).

**Figure 1.**
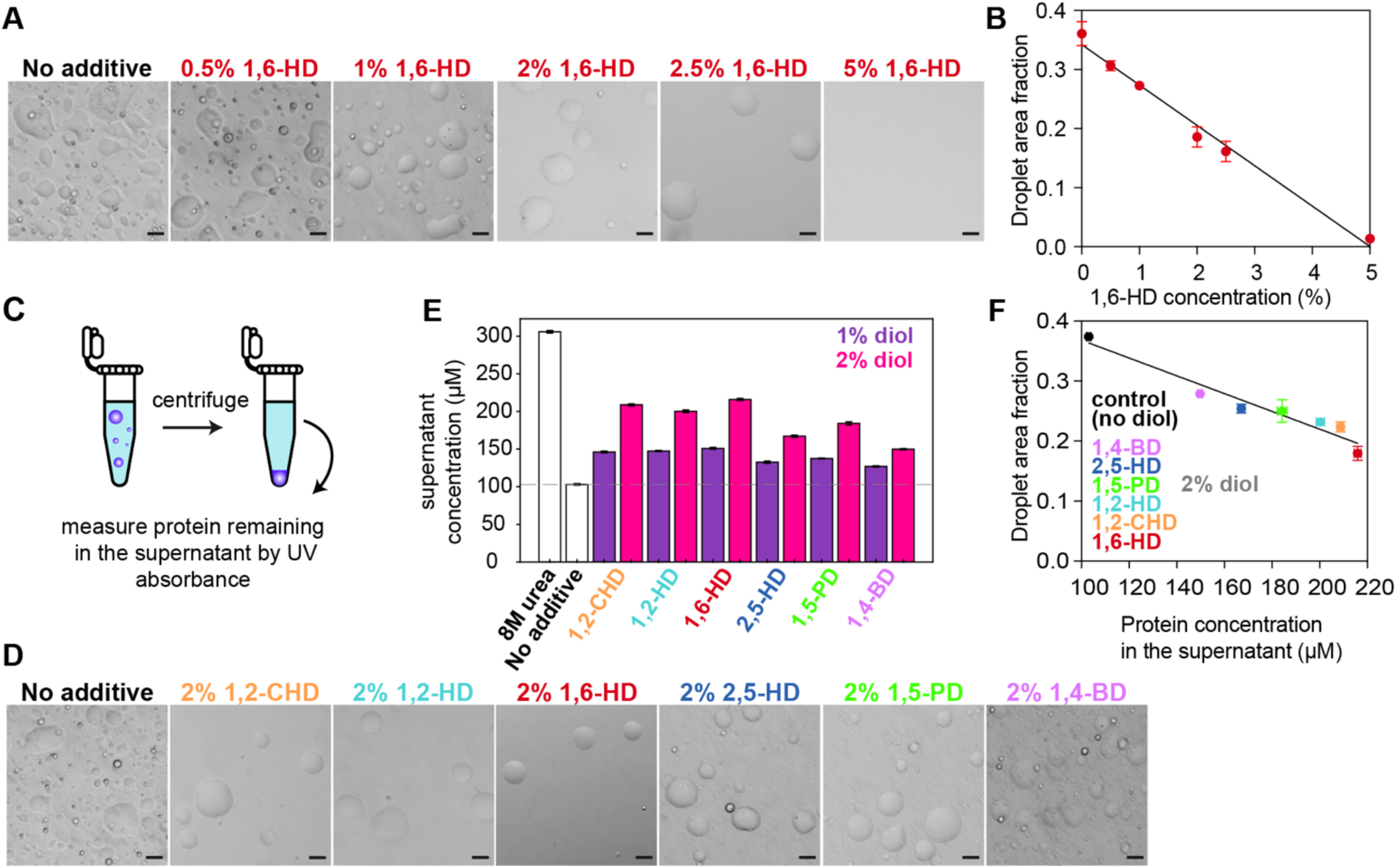
Alkanediols quantitatively alter FUS LC phase separation. A) DIC micrographs of 300 μΜ FUS LC in buffer alone (50 mM MES pH 5.5, 150 mM NaCl) and increasing concentrations of 1,6-hexanediol. Scale bars represent 20 μm. B) Extent of FUS LC phase separation in the presence of 0% to 5% 1,6-HD quantified by fluorescence microscopy. C) Schematic of experimental setup for quantifying phase separation measuring the concentration of protein remaining in the supernatant after centrifugation of droplets. D) DIC micrographs of 300 μM FUS LC in buffer alone (50 mM MES pH 5.5, 150 mM NaCl) (control) or various alkanediols with different aliphatic chain length or configuration premixed with buffer at 2%. Scale bars represent 20 μm. E) Phase separation assay that measures the saturation concentration of FUS LC in the presence of 1% or 2% of different alkanediols. F) Phase separation quantification using fluorescence microscopy versus supernatant concentration assay shows a high level of agreement between the two methods. Data in this figure are plotted as mean ± s.d. of n=3 technical replicates.

To compare the effect of various alkanediols, we imaged FUS LC droplets after treatment with an intermediate amount (2%) of each alkanediol (an amount suggested to be in the optimal range for cell-based experiments (Klein *et al*., 2020; Sabari *et al*, 2018; Shi *et al*, 2021)) using DIC as well as fluorescence (**Figure 1D, Figure S1D**). In separate samples, we quantified FUS LC phase separation by measuring *C*_sat_ in the presence or absence of each alkanediol at 1% or 2% concentration (**Figure 1E**). The impact on phase separation as measured by quantitative microscopy and *C*_sat_ measurements are highly correlated (**Figure 1F**), confirming the quantitative differences in disruption of FUS LC phase separation between the various alkanediols. Compared to the rest of the alkanediol series, 2,5-HD and 1,4-BD resulted in lower concentrations of protein remaining in the dispersed phase and higher droplet area fractions (**Figure 1F**), suggesting that these alkanediols were the least effective in disrupting the condensed phase. 1,5-PD reduced the extent of phase separation to an intermediate level, less than 1,6-HD and 1,2-HD. Together, the data suggest that alkanediols with different molecular structures all disrupt phase separation but to differing extents.

### Phase separation driven by charge-charge interactions is insensitive to alkanediols

In FUS, the N-terminal LC is followed by multiple domains involved in RNA-binding – the RNA-recognition motif (RRM) and zinc finger (ZnF) domains, flanked by interdomain linkers containing disordered RGG boxes composed of closely spaced arginine-glycine-glycine repeats and aromatic residues (e.g. RGG[Y/F]RGG) (Thandapani *et al*, 2013) that contribute to phase separation via protein-protein (Wang *et al*, 2018) and protein-RNA interactions (Chong *et al*., 2018). Previous studies have shown that the isolated RRM and ZnF domains of FUS bind RNA rather weakly, but the inclusion of RGG2 domain in RRM-RGG2 results in a many-fold increase in RNA-binding affinity, likely due to the presence of positively charged arginines that are well-known for promoting multivalency in disordered protein-RNA complexes (Loughlin *et al*., 2019; Ozdilek *et al*, 2017). Inspired by recent studies on the effect of hexanediol in the organization of RNA granules (Fuller *et al*, 2020) and chromatin (Itoh *et al*, 2021; Liu *et al*., 2021; Shi *et al*., 2021; Ulianov *et al*, 2021), as well as reports on the action of chemotherapeutics on nucleolar proteins and ribosomal RNA (rRNA) synthesis (Sutton & DeRose, 2021), we studied the potential of alkanediols to inhibit phase separation of FUS in the presence of RNA as a model for the types of protein-RNA interactions that contribute to biomolecular condensates formed in cells. As in previous studies (Burke *et al*, 2015; Monahan *et al*., 2017), we imaged full-length (FL) FUS after addition of TEV protease to cleave an N-terminal maltose binding protein (MBP) tag that prevents phase separation (**Figure 2A, S2A**). As expected, alkanediols disrupted the phase separation of FL FUS (**Figure 2A, 2B**). As we showed previously, addition of polyuridylic acid (polyU RNA) enhances FUS phase separation. However, we found that phase separation is only modestly decreased and still occurs even with the addition of 5% alkanediols, and the small differences in impacts on phase separation do not appear to correlate with the “strength” of the alkanediol in disrupting FUS phase separation without RNA. In other words, 1,6-HD and other alkanediols do not fully dissolve full-length FUS condensates with RNA.

**Figure 2.**
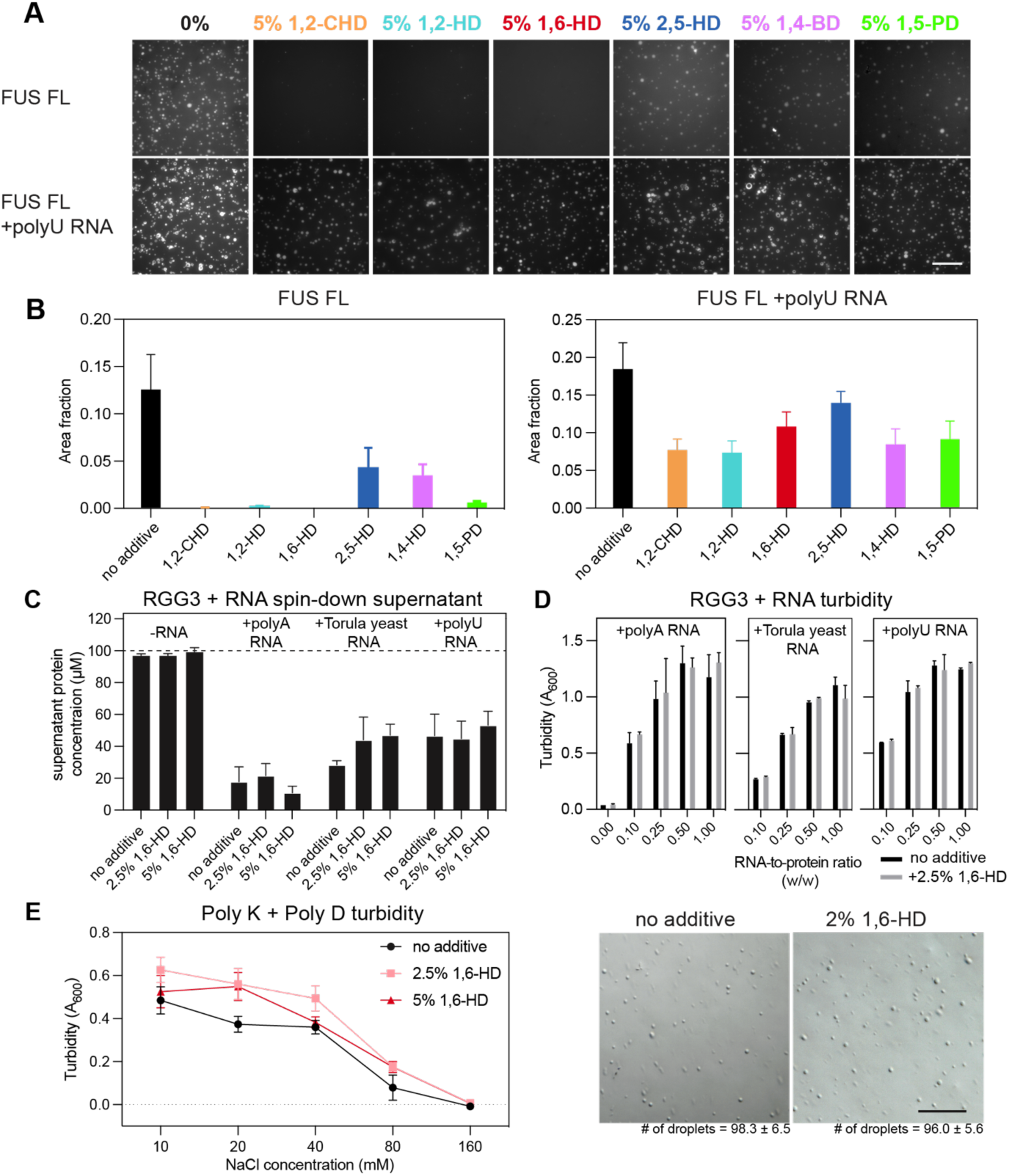
Alkanediols do not effectively disrupt charge-driven phase separation. A) Fluorescence microscopy images of 5 μΜ FUS full-length without (top) and with (bottom) Polyuridylic acid (0.05 mg/ml polyU, 5:1 protein-to-RNA ratio w/w) after treatment with 5% w/v premixed solutions of aliphatic alcohols and 20 μM ThT. Scale bars represent 50 μm. B) Phase separation of 5 μM FUS full-length without (left) and with (right) polyU RNA quantified by fluorescence-positive area fraction. Error bars represent standard deviation of three replicates. C) The effect of 2.5% or 5% 1,6-HD on phase separation of 100 μM FUS RGG3 in the presence of 1:1 (w/w) RNA measured by spin-down supernatant concentration. (Data are mean / standard deviation of three replicates.) D) The effect of 2.5% 1,6-HD on phase separation of 100 μM FUS RGG3 in the presence of RNAs across RNA-to-protein ratios, measured by turbidity at 600 nm wavelength. E) (left) The effect of 2.5% or 5% 1,6-HD on phase separation of the mixture of 2 mM poly-L-lysine and 2 mM poly-L-aspartate measured by turbidity at 600 nm. Error bars represent the standard deviation of three replicates. (right) DIC micrographs of 2mM poly-L-lysine and 2mM poly-L-aspartate mixture with and without 2% 1,6-HD added show similar droplet counts (average of triplicate) in a square with edge length 200 μm. Scale bars represent 50 μm.

Having already shown that RNA does not interact with FUS LC (Burke *et al*, 2015), we focused on an RGG domain of FUS as a model for the disordered, charged domains of FUS that may interact with RNA. FUS RGG3, which does not undergo phase separation by itself at these conditions, readily forms droplets upon the addition of RNA (**Figure S3**). Notably, the presence of alkanediols did not alter this behavior. For three different types of RNA (polyA, torula yeast RNA extract, and polyU) that include homopolymeric unstructured RNA and physiologically structured RNAs, we quantified the *C*_sat_ of FUS RGG3 in the phase-separated RGG3-RNA mixture by measuring the absorbance at both 260 and 280 nm to separate the contribution of protein and RNA to the readings. We found that *C*_sat_ values were nearly unchanged in the presence of 1,6-HD (**Figure 2C**). We also tested the impact of 2.5% 1,6-hexanediol on mixtures of FUS RGG3 and RNAs across various RNA-to-protein ratios and observed no significant disruption of phase separation at RNA-to-protein ratios up to 1:1 (**Figure 2D**).

Given that the co-phase separation of RGG3-RNA may rely predominantly on charge-charge interactions, we asked whether 1,6-HD is similarly incapable of disrupting other simpler complex coacervation driven by charged groups. To test this, we examined the phase separation behavior of poly-L-lysine (polyK) and poly-L-aspartate (polyD) peptide mixture (**Figure 2E**). As previously reported (Cakmak *et al*, 2020; Perry *et al*, 2015), the negatively charged polyD and positively charged polyK peptides readily phase separated upon mixing, and the turbidity of this peptide decreased sharply with increasing NaCl concentration, which screens charge-charge interactions between the peptides. Importantly, upon the addition of up to 5% 1,6-HD, we saw no apparent decrease in phase separation by microscopy; instead, the 1,6-HD additive appeared to slightly enhance turbidity, possibly due to the effect on charge screening of the decreased dielectric constant for solutions containing large volume fractions of alcohols like 1,6-HD.

Together, these findings suggest that while 1,6-hexanediol readily disrupts phase separation of FUS LC, it does not appear to disrupt coacervation involving RNA interaction with FUS RGG or oppositely charged peptide interactions.

### FUS LC chemical environment and molecular motion are altered by alkanediols

We then used NMR spectroscopy to obtain more detailed insight into how 1,6-HD interferes with FUS LC phase separation. Leveraging the uniquely sensitive, residue-by-residue resolution of ^1^H-^15^N heteronuclear single quantum coherence (HSQC) spectra (**Figure 3A**), we observed small chemical shift perturbations (CSPs) between the 0% and 5% 1,6-HD conditions throughout FUS LC domain (**Figure 3B**), that may arise due to weak interactions between 1,6-HD and the protein and/or small changes in the protein conformation or interactions with water. While the CSPs are smaller than those typically associated with strong binding interactions, they are larger than those observed for the weak interactions of karyopherin-β2 with FUS LC (Yoshizawa *et al*, 2018). The observed CSPs are also not uniform across the sequence. Binning backbone CSPs according to amino acid types showed little to no systematic residue-type specific effects and a broad distribution of changes within each type (**Figure 3C**). Consistent with the picture from the amide positions, we also observed small CSPs without clear residue-type specificity for backbone carbonyl (^13^CO) positions in the presence of 1,6-HD (**Figure S4**). Given the view that 1,6-HD is a hydrophobic disruptor, one might expect that hydrophobic amino acids in FUS may experience more chemical environment perturbation. However, neither these backbone CSPs above nor CSPs for sidechains from ^1^H-^13^C HSQC spectra (**Figure S4B**) showed large perturbations at hydrophobic amino acid positions. To examine if the range of 1,6-HD induced CSPs are specific to the FUS LC composition, we examined the backbone amide CSPs for FUS RGG3, which does not undergo phase separation at these concentrations and conditions and has a distinct sequence composition (Murthy *et al*., 2021). We observed a similar magnitude of CSPs for FUS RGG3 (**Figure S5**) as for FUS LC, suggesting that 1,6-HD similarly influences the chemical environment of distinct sequences. Therefore, our CSP analysis suggests that, despite certain residues showing larger CSPs, no clear pattern of interaction emerges.

**Figure 3.**
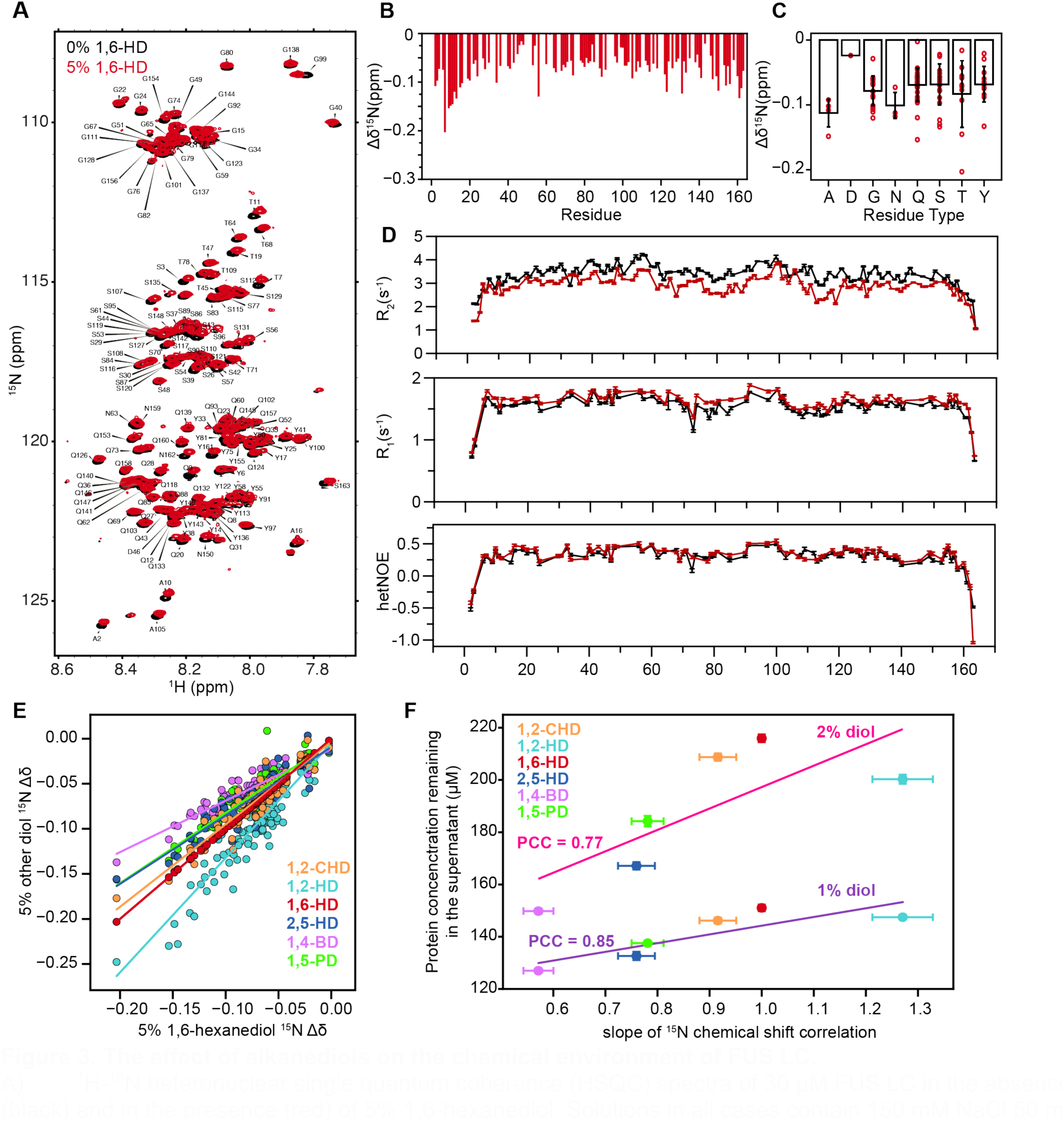
The effect of alkanediols on the chemical environment of FUS LC. A) ^1^H-^15^N heteronuclear single quantum coherence (HSQC) spectra of 30 μM FUS LC in the absence (black) and in the presence (red) of 5% 1,6-hexanediol. Solutions in all cases contain 150 mM NaCl 50 mM MES pH 5.5. B) FUS SYGQ LC ^15^N chemical shift deviations induced by 5% 1,6 hexanediol. Gaps indicate positions with no ^1^H/^15^N resonance (e.g. proline residues that have no backbone ^1^H attached to nitrogen) or with overlapped resonances that were not resolved in these experiments. C) 1,6-hexanediol induced ^15^N chemical shift deviations of FUS LC binned by residue type. Individual residues are plotted as red marks. Bar plots represent mean and standard deviation among all chemical shifts of each residue type. D) NMR spin relaxation parameters ^15^N *R*_2_, ^15^N *R*_1_ and (^1^H) ^15^N heteronuclear NOE values for FUS LC at 850 MHz ^1^H frequency indicate slightly faster molecular motions in the presence of 1,6-hexanediol. E) Comparison of ^15^N Δδ in 5% 1,6-hexanediol with every other co-solvent. F) Slope extracted from the correlation presented in (E) versus protein remaining in the supernatant shows that the chemical shift differences induced by each diol are correlated with the capacity of FUS LC to phase separate in each condition (PCC = 0.85 at 1% diol, PCC = 0.77 at 2% diol). Solid lines represent linear fits.

Next, we examined the changes in local molecular motions of FUS LC due to 1,6-HD by measuring the ^15^N NMR spin relaxation parameters (*R*_1_, *R*_2_, hetNOE) at each backbone amide position (**Figure 3D**). These values are sensitive to reorientational motions of the amide (NH) bond vector on the ps to ns timescales (Palmer *et al*, 2001). Decreased *R*_2_ and elevated *R*_1_ values in the presence of hexanediol suggest faster local reorientational tumbling motions, whereas the magnitude of the hetNOE (i.e. below 0.5) is unchanged and consistent with disorder across the domain, confirming that 1,6-HD does not cause significant structural rearrangements of the isolated LC domain. Interestingly, the decrease in ^15^N *R*_2_ observed for the addition of 1,6-HD to FUS LC is similar to that caused by introducing phosphomimetic serine-to-glutamate low complexity domain substitutions (FUS LC 12E) (**Figure S6A**), preventing phase separation (Monahan *et al*., 2017). Hence, elevated *R*_2_ values in wild-type FUS LC without 1,6-HD likely arise from transient favorable contacts between positions that also contribute to driving phase separation (Martin *et al*, 2020), and are suppressed by charged residue substitution (e.g. FUS LC 12E) or, as shown here, addition of 1,6-HD. Furthermore, addition of 1,6-HD to FUS LC 12E causes much smaller changes compared to the effects on FUS LC (**Figure S6B, S6C**), suggesting that 1,6-HD and the introduction of the 12 phosphomimetic mutations may weaken the same intramolecular contacts (**Figure S6**).

In an attempt to understand the mechanistic basis for the differing extents of phase separation disruption we observed for each alkanediols, we then sought to compare the effects of the different alkanediols on FUS LC by NMR. We acquired ^1^H-^15^N HSQC spectra of FUS LC domain in the presence of 0%, 2.5%, or 5% of each alkanediol and mapped the CSPs to identify locations with changes in chemical environment (e.g. conformational change or interaction) (**Figure S7**). Interestingly, when we plot each residue’s CSP upon addition of each alkanediol compared to those for 1,6-HD, we find a strong correlation (**Figure 3E, S8A,B**). Furthermore, the magnitude of the CSPs is strongly correlated with the *C*_sat_ of these molecules in the phase separation assay (**Figure 3F, S8C,D**), as shown by Pearson’s correlation coefficient (PCC = 0.85 at 1% diol, PCC = 0.77 at 2% diol for ^15^N CSPs, and PCC = 0.92 at 1% diol, PCC = 0.84 at 2% diol for ^1^H CSPs). Hence, the changes in residue-by-residue perturbation of the chemical environment are strongly correlated with the extent of phase separation disruption. In summary, we observe that the series of alkanediols show the same pattern of effects on FUS LC with different magnitudes that correlate with the extent of impact on phase separation. This observation suggests that all alkanediols, regardless of structure, have similar types of interactions with FUS. Hence, we do not find evidence here for unique interactions only possible by particular alkanediol isomers, though below we probe the factors that govern the extent of impact on phase separation.

### Alkanediols disrupt hydrophobic interactions

Structural differences (i.e., placement of the hydroxyl groups) have been suggested to make 2,5-HD less hydrophobic than 1,6-HD (**Figure 4A**), though the hydrophobicity of these and other alkanediols has, to our knowledge, never been experimentally measured. Using NMR-based concentration quantification experiments, we measured the partition coefficient (logP) between octanol and water, a common measure of hydrophobicity, for each alkanediol (**Figure 4B**) (Cheng *et al*, 2007). We find a range of partition coefficients showing that the longer alkanediols are more hydrophobic and the 2,5-HD isomer is indeed less hydrophobic than 1,6-HD. We then compared the measured logP with the slope of the NMR ^15^N chemical shift perturbations and with the *C*_sat_ of each diol from FUS LC phase separation assays (**Figure 4C**). We saw a strong correlation between the NMR shifts and logP, indicating that more hydrophobic alkanediols induce higher CSPs. Additionally, the correlation between *C*_sat_ and logP suggests that the hydrophobicity of the diols is linked to their ability to disrupt phase separation. However, 2,5-HD and 1,2-HD are outliers in the correlation between *C*_sat_ and logP where removing these outliers significantly improves the correlation (PCC 0.76 to 0.99). This suggests that although hydrophobicity plays the dominant role, the unique geometries of these branched alkanediols do introduce additional contributions to their impact on phase separation, making them different from the alkanediols with hydroxyl groups at the terminal positions.

**Figure 4.**
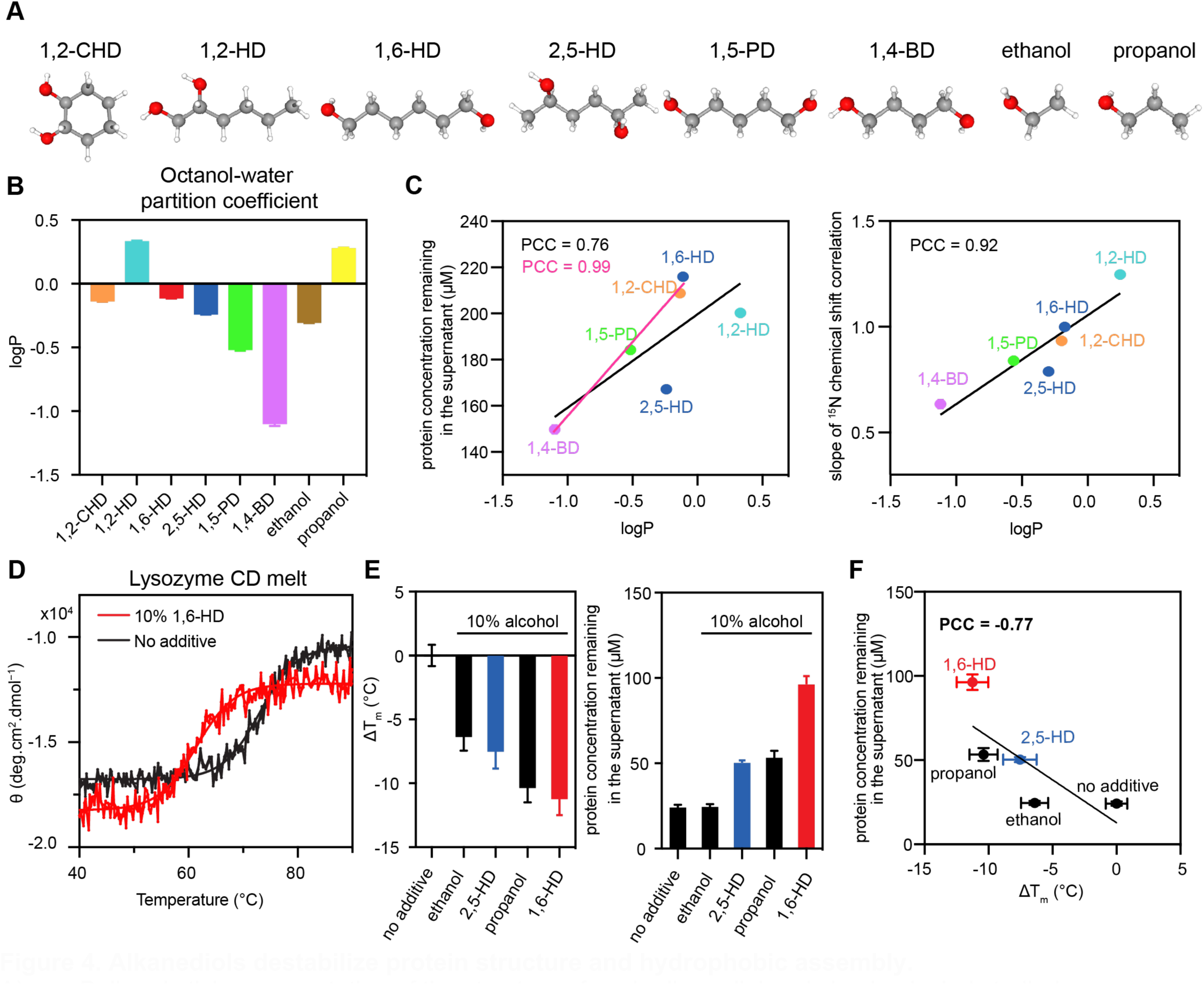
Alkanediols destabilize protein structure and hydrophobic assembly. A) Ball and stick representation of the structure of each alkanediol and simple alcohol studied. B) NMR measured values of the octanol partition coefficient of each alkanediol, a measure of hydrophobicity. Log P corresponds to the logarithm of the ratio of the concentration of the solute in octanol versus in water. C) Correlation of logP presented in (B) versus ^15^N Δδ slopes in Figure 3F (top), or versus protein remaining in the supernatant presented in Figure 1D (bottom) shows that the effect of the alkanediols on protein backbone chemical shifts and FUS phase separation is correlated with the hydrophobicity. Correlation with all data points is shown in black, while the correlation excluding 2,5-HD and 1,2-HD is indicated in magenta. D) Circular Dichroism data showing the effect of 10% 1,6-HD on the thermostability of lysozyme. E) (left) The melting temperature of lysozyme decreases with the addition of simple alcohols as well as diols. (right) Saturation concentration of FUS LC-RGG with and without the addition of simple alcohols or diols. Error bars represent the standard deviation of three replicates. F) Correlation between saturation concentration of FUS LC-RGG and melting temperature change (ΔTm) of lysozyme with different alcohols.

Taking inspiration from prior studies that have established the capability of aliphatic alcohols such as ethanol and propanol to destabilize protein structures by weakening hydrophobic interaction (Chong *et al*, 2015; Miyawaki & Tatsuno, 2011; Nakata *et al*, 2023), we sought to determine whether 1,6-HD, like its simpler counterparts, could exert a similar influence on hydrophobic interactions and protein structures. Specifically, we used circular dichroism as a function of temperature to assess the impact of 1,6-HD on the thermal stability of lysozyme—a protein that folds around a hydrophobic core (**Figure 4D**). Interestingly, 10% 1,6-HD reduced the melting temperature (T_m_) of lysozyme by approximately 10 °C. Next, we expanded our analysis to include 2,5-HD and other representative simple alcohols, such as ethanol and 1-propanol. We found that the extent of lysozyme T_m_ reduction for 1,6 HD is comparable to propanol, which has the same number of linear aliphatic carbons per hydroxyl group (**Figure 4E**), while 2,5-HD and ethanol have less impact on the T_m_. Furthermore, we compared the structure-destabilizing capacity of the alcohols to their effect on FUS phase separation (**Figure 4E,F**). For these experiments, we used a longer segment, FUS LC-RGG1 that we recently characterized, due to its increased ability to undergo phase separation compared to FUS LC (Wake *et al*., 2024), allowing quantification of the impact of 10% 1,6-HD and matching those we used in the CD studies. Interestingly, the lysozyme T_m_ reduction of each alcohol correlates moderately (PCC = 0.77) with their individual potency in reducing phase separation (**Figure 4F**). This observation suggests a shared mechanism for alcohol-induced reduction in phase separation and folding stability. Among all molecules tested, those with longer aliphatic chains (i.e., propanol and 1,6-HD) exhibited greater potency in disrupting lysozyme fold and FUS LC-RGG1 phase separation, again suggesting that the hydrophobicity of the alcohol plays a pivotal role in destabilizing both folding and phase separation.

### 1,6-Hexanediol perturbs protein-protein interactions via direct contacts with protein

To gain more atomistic details about the impact of alcohols on FUS LC phase separation, we performed atomistic MD simulations of an isolated single chain of FUS LC solvated with water and ion molecules (“in explicit solvent”) in the presence of 5% 1,6-HD or 2,5-HD and compared these simulations to those without hexanediols. The parameters for both hexanediols, modeled with GAFF2 force field (He *et al*, 2020) (see **Methods** for more details), were validated by comparing the densities and molar volumes of water solutions with experimental data (Romero *et al*, 2007) (**Figure S9** and **Appendix Tables S1** and **S2**).

Analysis of the protein conformation, including the radius of gyration (R_g_) distribution and the average intrachain distance (R_ij_), revealed that both alkanediols induced a more expanded structure for the LC domain compared to simulations without them (**Figure 5A and inset**). Additionally, the expansion was more pronounced with 1,6-HD compared to 2,5-HD, which is consistent with 1,6-HD’s stronger potency in disrupting FUS LC phase separation. Notably, FUS LC with 1,6-HD exhibited the largest R_ij_ expansion, especially for residues that are furthest apart (| i – j | > 100), consistent with disruption of long-range protein-protein interactions. Contact analysis (see Methods) further supported these findings, showing a decrease in intramolecular protein-protein contacts in presence of either alkanediols, with a greater reduction observed with 1,6-HD (**Figure 5B**). Particularly, 1,6-HD significantly disrupted mid- to long-range contacts, as indicated by a region of reduced values near the top of the contact heat map, as well as the distribution of average protein-protein contacts across various ranges of residue distances (**Figure 5C**), which aligns with the observed R_ij_ expansion shown in **Figure 5A**.

**Figure 5:**
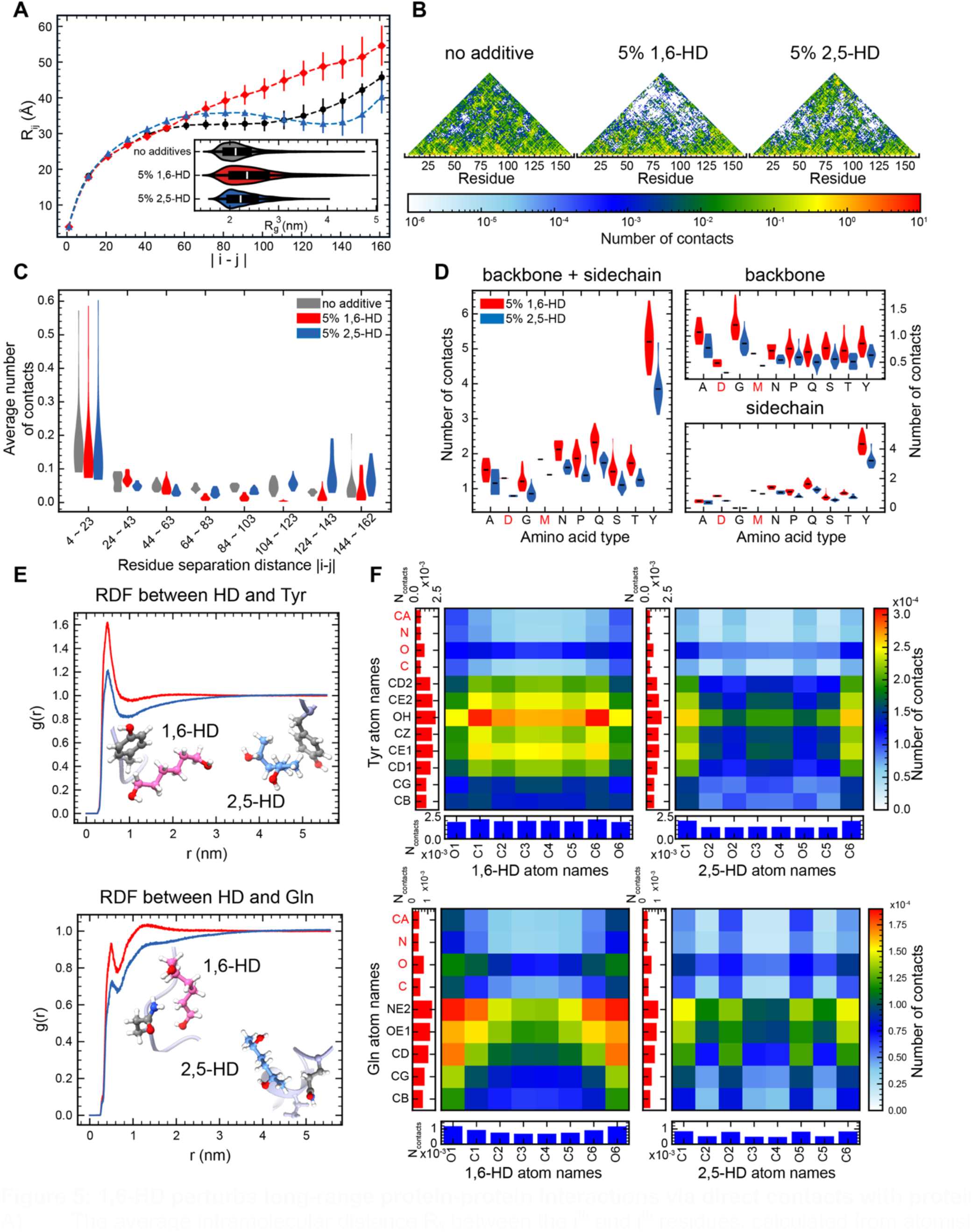
1,6-HD perturbs long-range protein-protein interactions via direct contacts with protein. A) The average intramolecular distance R_ij_ between the i^th^ and j^th^ residues, calculated from atomistic simulations of single chains FUS LC with 100 mM NaCl without additives, with 5% 1,6-hexanediol (5% 1,6-HD), or with 5% 2,5-hexanediol (5% 2,5-HD). Standard error of the mean is computed over 3 independent trajectories for each condition. Inset: Distribution of radius of gyration (R_g_) calculated from the simulations. B) Protein-protein contact profiles calculated from the simulations. From left to right show the result from no additive, 5% 1,6-HD, and 5% 2,5-HD simulations, respectively. C) The distribution of average protein-protein contacts in different ranges of residue distances. D) The distribution of protein-hexanediol contacts calculated from the simulations, binned by residue type. E) Radial distribution functions (RDF, g(r)) between the center of mass (COM) of sidechain heavy atoms of Tyr / Gln and heavy atoms of hexanediols. Corresponding interaction example snapshots are inset. F) Atomistic contact maps of Tyr / Gln with hexanediols, calculated from the atomistic simulations. The left column and the bottom row of each panel show the one-dimensional summation of corresponding atomistic contacts.

Previous research has indicated that small molecules can influence proteins through direct interactions (Das & Mukhopadhyay, 2009; Nakata *et al*., 2023). To explore the potential disruptive mechanism of hexanediol, we investigated protein-alkanediol contacts in our simulations, which revealed extensive contacts formed between alkanediols and the entire protein chain (**Figure 5D** and **S10A**), consistent with NMR data showing widespread CSPs among the protein. Notably, 1,6-HD formed more contacts with protein than 2,5-HD. Moreover, the radial distribution functions (RDF, g(r)) for protein residues with hexanediols showed more close contact formation and surface enrichment for 1,6-HD (**Figure 5E** and **S10B**). These findings suggest stronger interactions between 1,6-HD and protein, consistent with its stronger potency in disrupting FUS phase separation. Further details from residue-diol contact maps show that hexanediol forms contacts with different residues using different parts of the molecule (**Figure 5F** and **S10C**). 1,6-hexanediol formed more contacts with Tyr using the aliphatic region and with Gln using the polar ends. Interestingly, the geometric differences between 1,6-HD and 2,5-HD lead to differences in the number of protein contacts. Complementarily, comparison of 1,6-HD and 2,5-HD NMR ^1^H-^13^C HSQC spectra showed that the middle aliphatic methylene group carbon of 1,6-HD was more shielded (lower chemical shift in ppm) than that of 2,5-HD (**Figure S11**), suggesting a less polar, more hydrophobic chain, which aligns with 1,6-HD’s ability to better interact with hydrophobic amino acid side chains. Notably, the central section of 2,5-HD also shows lower contact frequency with Tyr than that of 1,6-HD (**Figure 5F**), which may be due to a combination of differences in steric hindrance, polarity, and hydrophobicity. In other words, the linear geometry of 1,6-HD appears to facilitate contacts with protein residues, in contrast to the branched heavy atom chain of 2,5-HD. Taken together, these data suggest that direct interaction between protein residues and hexanediols can contribute to phase separation disruption and that some of the difference between 1,6-HD and 2,5-HD can be explained by changes in contacts caused by their distinct molecular geometries as proposed previously (Gu *et al*, 2023).

## Discussion

In this study, we demonstrated that alkanediols of different lengths and configurations, including 1,4-BD, 1,5-PD, 1,2-HD, 1,2-CHD, 1,6-HD, and 2,5-HD, can effectively decrease phase separation of FUS. Although very high (60%) hexanediol concentrations has been suggested to cause the ordering of an unstructured region in a protein crystal (Buhrman *et al*, 2003), our data suggest no change to the overall disordered structure of FUS LC in the concentration range (up to 10%) used in cellular studies. In fact, faster reorientational motions for FUS LC with 1,6-HD likely stem from decreased self-interaction within FUS molecules. Among tested compounds, 1,6-HD stands out as the most effective disruptor of FUS phase separation. Indeed, 2,5-hexanediol is less effective than 1,6-hexanediol in dissolving nuclear speckles and subsequent recruitment of a proline-glutamine-rich splicing factor (Levone *et al*, 2021) and melting FUS LC hydrogels and intracellular puncta (Kato & McKnight, 2018), and it shows significantly less impact on FUS phase separation (**Figures 1 and 3**). However, we show here that despite being less potent than 1,6-HD, 2,5-HD and shorter alkanediols are still able to reduce phase separation via a qualitatively similar mechanism. Taken together, these results suggest that specific interactions uniquely available to particular alkanediol isomers (e.g. simultaneous hydrogen bonding of both hydroxyl groups only geometrically possible in 1,6-HD and not 2,5-HD) do not explain the differences in their action. Rather, on the atomic level, the aliphatic segment of 1,6-HD allows interactions with hydrophobic and aromatic amino acids such as tyrosine, while the polar hydroxyl groups enable hydrogen bonds with polar residues such as glutamine, suggesting it may compete with both of these residue types for protein-protein contacts that drive phase separation (Murthy *et al*., 2019; Wake *et al*., 2024; Wang *et al*., 2018). Furthermore, these simulations and experiments do provide insight into how hydrophobicity and geometry contribute to the differences in potency between 1,6-HD and 2,5-HD. Specifically, the linear aliphatic chain present in 1,6-HD is less polar and more accessible for interaction with hydrophobic protein residues, while the primary alcohol in 1,6-HD is more polar than the secondary alcohol in 2,5-HD, hence more favorable for dipole-dipole interactions. Both factors combine to make 1,6-HD more amphiphilic than 2,5-HD, consistent with 1,6-HD’s greater ability to suppress phase separation than 2,5-HD.

Consistent with earlier biophysical studies on FG-rich nucleoporins and the synaptonemal complex (SC) that proposed that the capacity of aliphatic alcohols to dissolve the NPC and SC is correlated with the hydrophobicity of each alkanediol (Ribbeck & Gorlich, 2002; Rog *et al*, 2017), we measured the hydrophobicity of each alkanediol and found that it correlates extremely well with the ability of alkanediols with terminal hydroxyl groups (1,6-HD, 1,5-PD, 1,4-BD) to disrupt phase separation. However, the unique structures of 1,2-HD and 2,5-HD result in less disruption in phase separation than predicted by hydrophobicity alone, suggesting that the geometry of the molecule does play a role in determining the capacity of the alcohols to disrupt FUS LC phase separation. Consistent with disruption of the hydrophobic driving forces for folding, we also show that aliphatic alcohols, including 1,6-HD, can also disrupt protein folding stability, suggesting that alkanediols may also unfold cellular proteins. Intriguingly, 1,2-HD also disrupted TEV protease activity (**Figure S2A**), complementing previous findings that 1,6-HD disrupts kinase and phosphatase activity (Düster *et al*, 2021). Therefore, our data suggest that disruptions beyond only disordered domain contacts may contribute to cellular changes caused by hexanediol treatment, and the mechanism of these changes should be inferred cautiously. Nevertheless, our biochemical data suggest that interference with hydrophobic interactions constitutes a major mechanism underpinning the disruptive function of 1,6-HD in the context of biochemical FUS phase separation, as also suggested by the impact of salts on the Hofmeister series on FUS phase separation (Murthy *et al*., 2019).

Some RNA-rich condensates in cells show insensitivity to 1,6-HD treatment (Ahmed *et al*, 2021; Sato *et al*, 2022). Our data show that the alkanediol-sensitivity of FUS condensates enriched in RNA is reduced compared to FUS without RNA (**Figure 2A**), and FUS RGG-RNA and polyK-polyD complex coacervation are largely unaffected by hexanediol treatment, suggesting that charge-charge interactions mediating phase separation are not impacted by amphiphilic alcohols. Although hexanediol-dependent dissolution has been successfully used to distinguish solid from liquid forms of particular membraneless organelles, such as those stabilized by hydrophobic interactions (Kroschwald *et al*., 2017), the fact that certain condensates are not susceptible to 1,6-hexanediol does not necessarily mean they are solid. Instead, these condensates may primarily involve charge-charge interactions and may indeed be liquid. As a corollary, the large number of membraneless organelles found to be susceptible to 1,6-hexanediol may also therefore imply that these are primarily stabilized by hydrophobic and not charge-charge interactions. However, these observations that interactions between disordered RGG peptides and unstructured RNA or total RNA extracts are not disrupted by 1,6-HD should not be taken to mean that no cellular RNA-protein interactions are disrupted by 1,6-HD. Indeed, FUS and many other phase-separating RNA-binding proteins contain RRMs and other domains that interact with single stranded RNA not primarily via the charged backbone of RNA but rather via the bases using hydrophobic/aromatic residues (Loughlin *et al*., 2019; Ozdilek *et al*., 2017; Ozguney *et al*, 2024), which may be disrupted by 1,6-HD. Thus, the effectiveness of hexanediol is context-dependent and results of cellular experiments adding HD should be interpreted with careful consideration of the condensate composition (e.g. sequence characteristics and nucleic acid partitioning).

Given the expanding knowledge of the roles of biomolecular condensates in many physiological processes (Lyon *et al*, 2021) and in diseases (Alberti & Dormann, 2019; Alberti & Hyman, 2021; Jiang *et al*, 2020; Lu *et al*, 2021), many efforts have focused on understanding how to modulate phase separation and how potential small molecule therapeutics partition into condensates (Dai *et al*, 2021; Klein *et al*., 2020; Wheeler *et al*., 2016). For example, mitoxantrone and other chemotherapy drugs with targets that reside in nuclear condensates selectively concentrate in nuclear condensates (Chong *et al*., 2015). This selective concentration may be due to direct interactions between the compound and the disordered protein – although these direct interactions are weak, they can show similarly sized NMR chemical shift perturbations at 3 orders of magnitude lower concentrations (500 μM vs ~400 mM) (Uechi *et al*, 2019) than perturbations generated by alkanediols (**Figure 3**), suggesting that small molecules other than hexanediol may be able to alter condensate properties. Our results show that NMR-based observations combined with molecular simulation can provide unique insight into the molecular origins of phase separation modulation and could contribute to the rational design of possible therapies for altering disordered protein phase separation.

## Supporting information

Supplemental Figurs and Table

## Acknowledgments

We thank Dr. Mandar Naik for NMR assistance and the Structural Biology Core Facility at Brown University. Research was supported in part by NIGMS R01GM147677 (to N.L.F.), Human Frontier Science Program RGP0045/2018 (to N.L.F), grant 1845734 from the National Science Foundation (to N.L.F.), and NIGMS R35GM153388 (to J. M.). T.Z. was supported in part by a Pape Adams Postdoctoral Award from the Carney Institute for Brain Science at Brown University and a Milton Safenowitz Postdoctoral Fellowship from the ALS Association. N.W. was supported in part by an NIGMS training grant at Brown University (T32GM139793). A.C.M. was supported in part by NIGMS training grant at Brown University (T32GM007601) and NSF graduate fellowship (1644760, to A.C.M.). This content solely reflects the authors and does not necessarily represent the official views of the funding agencies. We gratefully acknowledge the computational resources provided by the Texas A&M High Performance Research Computing (HPRC).

## Author contributions

T.Z., N.W., T.M.P., A.C.M., R.W., and N.L.F. designed experiments for NMR spectroscopy, microscopy, and phase separation assays. T.Z. and T.M.P. performed and analyzed data for NMR spectroscopy, N.W. and T.M.P. performed microscopy and phase separation assays. T.Z., T.M.P., and N.L.F wrote the manuscript with contributions from all authors. S.L.W. and J.M. designed, performed, and analyzed data for MD simulations.

## Experimental procedures

### General Information

1,2-hexanediol (#213691), 1,6-cyclohexanediol (#141712), 1,6-hexanediol (#240117), 2,5-hexanediol (#H11904), 1,4-butanediol (#493732) and 1,5-pentanediol (#P7703) were purchased from Sigma Aldrich.

### Protein Purification

FUS LC containing a TEV cleavable N-terminal histidine tag (RP1B FUS LC, AddGene #127192), full-length FUS with a TEV cleavable N-terminal histidine / maltose-binding protein (MBP) fusion tag (pTHMT FUS 1-526, AddGene #98651), FUS LC-RGG1 containing a TEV cleavable N-terminal histidine tag (pRP1B FUS LC-RGG1 (1-284)) and FUS RGG3 with a TEV cleavable N-terminal histidine / MBP fusion tag (pTHMT FUS RGG3 (453-507)) were expressed in BL21 Star (DE3) *Escherichia coli* cells (Life Technologies) grown at 37°C to an OD of 0.60-0.90 before induction with 1 mM IPTG for 4 hours. Isotopically labeled protein was produced by expression in M9 minimal media supplemented with ^15^N-ammonium chloride or ^13^C-glucose (Cambridge Isotopes). Histidine tagged FUS LC and FUS LC-RGG1 was purified as described previously (Wake et al, 2024). In brief, cells were lysed using an Avestin homogenizer and the lysate was cleared by centrifugation at 20,000 rpm for 1 hour. The insoluble fraction was applied to a 5 mL HisTrap HP column (Cytiva) equilibrated with 8 M urea 20 mM NaPi pH 7.4 300 mM NaCl 10 mM imidazole and eluted with a gradient from 10-300 mM imidazole. The protein was diluted with 20 mM NaPi pH 7.4 such that the final urea concentration was 1 M and incubated with TEV protease over-night. A subtractive nickel affinity step was performed, and the protein in the flow through was then buffer exchanged into 20 mM CAPS pH 11.0 and concentrated for storage by centrifugal filtration. MBP-FUS 1-526 was purified as previously described (Burke *et al*, 2015). MBP-FUS RGG3 was purified as previously described (Murthy et al, 2021). All protein preparations showed UV absorbance ratio, A_260 nm_/A_280 nm_, of 0.5 to 0.6, indicating minimal to no residual nucleic acid contamination. Proteins were stored as frozen aliquots at −80 °C.

### Phase separation assays

Phase separation of FUS LC in the presence of hexanediols was quantified by measuring the absorbance at 280 nm of the dilute phase using a NanoDrop spectrophotometer. Samples were prepared by diluting FUS LC stored in 20 mM CAPS pH 11.0 to a final protein concentration of 300 μM into 20 mM MES (pH adjusted with Bis-Tris) pH 5.5 150 mM NaCl with and without hexanediols. After dilution, samples were spun at 14,000 g for 10 minutes at 22°C. Phase separation of FUS LC-RGG in the presence of hexanediols was quantified the same way as FUS LC.

Turbidity of RGG3 was measured as previously described (Murthy et al, 2021) by measuring the absorbance at 600 nm of samples in a 96-well clear plate (Costar) using a Cytation 5 Cell Imaging Multi-Mode Reader (BioTek). Samples were prepared in 50 mM MES pH 5.5, 150 mM NaCl buffer. For turbidity of FUS FL, TEV protease was added to 10 μM MBP-FUS FL in 50 mM HEPES pH 7 with 150 mM NaCl and, after 60 minutes, buffer containing 10% alkanediols and RNA, for a final concentration of 5% alkanediol. To remove background absorbance, the turbidity of a no TEV control (i.e. with TEV storage buffer) for each condition was subtracted from the turbidity of the experimental conditions. Experiments were conducted in triplicate and averaged. To test the effect of different alkanediols on RNA-FUS phase separation, the polyadenylic acid (polyA), polyuridylic acid (polyU), and torula yeast RNA extract type VI (Sigma #R6625) were desalted into the appropriate buffer using a Zeba 0.5 ml spin column.

To test the effect of 1,6-HD on polyD + polyK phase separation, the peptides were dissolved in 50 μM MES pH 5.5 buffer, turbidity of 2 mM polyK + 2 mM polyD mixtures with 10-160 mM NaCl was measured in a 96-well clear plate (Costar) using the Cytation 5 reader by measuring the absorbance at 600 nm. Experiments were conducted in triplicate and averaged. Poly-L-lysine hydrochloride (PLKC10, #26124) and poly-L-aspartic acid sodium salt (PLD10, #34345) were purchased from Alamanda Polymers.

### Microscopy

Visualization of phase separation of FUS LC, MBP-FUS FL and FUS RGG3 was performed by differential interference contrast microscopy on a Zeiss Axiovert 200M Fluorescence microscope. All samples were spotted on a 24×40×1.5 mm coverslip for imaging using a 40x water immersion microscope objective. Images were processed using ImageJ (NIH). Fluorescence microscopy samples were prepared as 200 μM FUS LC or (60 μM FUS FL) in a 50 mM HEPES buffer (pH adjusted to 7.0 using Bis-Tris) containing 150 mM sodium chloride, 20 μM Thioflavin T as a fluorescent dye, and varying volume fractions of alkanediols. All samples were spotted onto the coverslip and droplets were allowed to settle for 2 minutes prior to capturing four independent fields of view for quantification. The droplet area ratio was determined using an in-house Matlab (2023b) script that calculates the percentage of fluorescently illuminated pixels corresponding to droplet area. Droplet counting for the polyK-polyD droplets were performed using ImageJ.

### NMR spectroscopy

NMR experiments were recorded at 850 MHz using a Bruker Avance III spectrometer with a ^1^H/^15^N/^13^C TCl cryoprobe and *z* field gradient coil. NMR titrations of ^15^N-labeled FUS LC with 0, 2.5 or 5% hexanediols were conducted at 25°C in 50 mM MES, 150 mM NaCl pH 5.5 including 10% ^2^H_2_O and 2 μΜ sodium trimethylsilylpropane sulfonate (DSS). All data were processed with NMRPipe software package (Delaglio *et al*, 1995) and visualized with NMRFAM-Sparky(Lee *et al*, 2015) or CcpNMR Analysis (Vranken *et al*, 2005). Chemical shifts and intensity ratios were normalized by subtracting the ^15^N chemical shift values and dividing the signal intensity in the absence of hexanediol from all other datasets. DSS was used as the direct reference for the ^1^H chemical shift, while the ^13^C and ^15^N chemical shifts were referenced indirectly using DSS using their gyromagnetic ratios. Specifically, adjustments in the nitrogen and carbon dimensions were calculated based on the following formula (Harris *et al*, 2001):

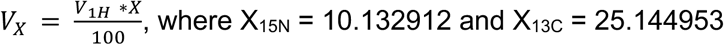

Molecular motions in the presence of 1,6-hexanediol were probed using ^15^N *R*_1_, ^15^N *R*_2_ and heteronuclear NOE experiments using standard pulse sequences (hsqct2etf3gpsitc3d, hsqct1etf3gpsitc3d, hsqcnoef3gpsi, respectively from Bruker Topspin 3.2). Interleaved experiments comprised 256 x 4096 total points in the indirect ^15^N and direct ^1^H dimensions, respectively, with corresponding acquisition times of 74 ms and 229 ms, sweep width of 20 ppm and 10.5 ppm, centered at 117 ppm and 4.7 ppm, respectively. ^15^N *R*_2_ experiments had an interscan delay of 2.5 s, a Carr-Purcell-Meiboom-Gill (CPMG) field of 556 Hz, and total *R*_2_ relaxation CMPG loop-lengths of 16.5 ms, 264.4 ms, 181.8 ms, 33.1 ms, 115.7 ms, 82.6 ms, and 165.3 ms. ^15^N *R*_1_ experiments had an interscan delay of 1.2 s, and total *R*_1_ relaxation loop-lengths of 100 ms, 1000 ms, 200 ms, 800 ms, 300 ms, 600 ms, and 400 ms. Heteronuclear NOE experiments were conducted with an interscan delay of 5 s.

### Circular Dichroism spectrometry of lysozyme

0.1 mg/mL lysozyme (Sigma #L4919) solution was prepared in 5 mM sodium phosphate buffer at pH 7. The solution was then filtered through a 0.22 μm membrane filter. Circular dichroism (CD) spectra of lysozyme were recorded using a JASCO J-815 spectropolarimeter equipped with a Peltier temperature control system. Molar residue ellipticity (MRE) at 222 nm was monitored in a temperature range between 25-90°C both in the presence and absence of 10% alcohol additives.

### Single chain MD simulations of FUS LC with or without hexanediol additives

Force field parameters for 1,6- and 2,5-hexanediol were obtained from the open source ACPYPE python package Antechamber (Sousa da Silva & Vranken, 2012) with the GAFF2 force field (He *et al*., 2020).

Initial equilibration simulations were performed using the classical MD package GROMACS-2022 (Abraham *et al*, 2015), employing periodic boundary conditions and a 2 fs integration time step. All systems were simulated in explicit solvent, modeled using the AMBER03ws force field and explicit TIP4P/2005s water model (Best *et al*, 2014) with improved salt (NaCl) parameters (Luo & Roux, 2009).

FUS LC initial protein structure was placed in a 12×12×12 nm octahedral box, and 330 hexanediol molecules were added to achieve a concentration of ~5%. The system was then solvated, with counter ions added to achieve electroneutrality and a salt concentration of 100 mM. It then underwent energy minimization using the steepest descent algorithm, followed by a 100 ps NVT equilibration. Velocity rescaling algorithm (Bussi *et al*, 2007) was used for temperature equilibration at 300 K with a coupling constant of 0.1 ps. Subsequently, a 100 ps NPT equilibration was conducted using the Parrinello-Rahman barostat (Parrinello & Rahman, 1980) with a coupling constant of 2 ps for pressure control.

The production runs were performed with MD package Amber22 (Case *et al*, 2022). After GROMACS equilibration, the package ParmEd (Shirts *et al*, 2017) was used to convert GROMACS files to Amber files. Hydrogen mass repartitioning (Hopkins *et al*, 2015) was applied during conversion to enable a timestep of 4 fs for production runs. Following the conversion, energy minimization was applied with protein position restraint, followed by a 2 ns NVT equilibration with reduced protein position restraint. With Berendsen barostat (Berendsen *et al*, 1984), a 500 ps NPT equilibration with further reduced protein position restraint was conducted. After the equilibration steps, the production run simulations were performed in the NPT ensemble. The Langevin dynamics (Pastor, 1994) were used to control temperature at 300 K (friction coefficient = 1 ps⁻¹), the Monte Carlo barostat (Åqvist *et al*, 2004) was applied to control pressure at 1 bar (coupling constant = 1), and the SHAKE algorithm, as implemented in Amber22, was used for constraining hydrogen-containing bonds.

After production runs, the Amber NetCDF trajectory files were converted to Gromacs compressed trajectory files via CPPTRAJ (Roe & Cheatham, 2013) package. Python package MDAnalysis-2.5.0 (Gowers *et al*, 2016) was used for further analysis.

## References

Abraham MJ, Murtola T, Schulz R, Páll S, Smith JC, Hess B, Lindahl E (2015) GROMACS: High performance molecular simulations through multi-level parallelism from laptops to supercomputers. SoftwareX 1-2: 19–25

Ahmed J, Meszaros A, Lazar T, Tompa P (2021) DNA-binding domain as the minimal region driving RNA-dependent liquid–liquid phase separation of androgen receptor. Protein Science 30: 1380–1392

Alberti S, Dormann D (2019) Liquid-Liquid Phase Separation in Disease. Annu Rev Genet 53: 171–194

Alberti S, Hyman AA (2021) Biomolecular condensates at the nexus of cellular stress, protein aggregation disease and ageing. Nat Rev Mol Cell Biol 22: 196–213

Altmeyer M, Neelsen KJ, Teloni F, Pozdnyakova I, Pellegrino S, Grofte M, Rask MD, Streicher W, Jungmichel S, Nielsen ML et al (2015) Liquid demixing of intrinsically disordered proteins is seeded by poly(ADP-ribose). Nat Commun 6: 8088

Åqvist J, Wennerström P, Nervall M, Bjelic S, Brandsdal BO (2004) Molecular dynamics simulations of water and biomolecules with a Monte Carlo constant pressure algorithm. Chemical Physics Letters 384: 288–294

Babinchak WM, Dumm BK, Venus S, Boyko S, Putnam AA, Jankowsky E, Surewicz WK (2020) Small molecules as potent biphasic modulators of protein liquid-liquid phase separation. Nat Commun 11: 5574

Banani SF, Lee HO, Hyman AA, Rosen MK (2017) Biomolecular condensates: organizers of cellular biochemistry. Nat Rev Mol Cell Biol 18: 285–298

Berendsen HJC, Postma JPM, Vangunsteren WF, Dinola A, Haak JR (1984) Molecular-Dynamics with Coupling to an External Bath. Journal of Chemical Physics 81: 3684–3690

Berkeley RF, Kashefi M, Debelouchina GT (2021) Real-time observation of structure and dynamics during the liquid-to-solid transition of FUS LC. Biophys J 120: 1276–1287

Best RB, Zheng W, Mittal J (2014) Balanced Protein-Water Interactions Improve Properties of Disordered Proteins and Non-Specific Protein Association. J Chem Theory Comput 10: 5113–5124

Bock AS, Murthy AC, Tang WS, Jovic N, Shewmaker F, Mittal J, Fawzi NL (2021) N-terminal acetylation modestly enhances phase separation and reduces aggregation of the low-complexity domain of RNA-binding protein fused in sarcoma. Protein Sci 30: 1337–1349

Boija A, Klein IA, Young RA (2021) Biomolecular Condensates and Cancer. Cancer Cell 39: 174–192

Buhrman G, de Serrano V, Mattos C (2003) Organic solvents order the dynamic switch II in Ras crystals. Structure 11: 747–751

Burke KA, Janke AM, Rhine CL, Fawzi NL (2015) Residue-by-Residue View of In Vitro FUS Granules that Bind the C-Terminal Domain of RNA Polymerase II. Mol Cell 60: 231–241

Bussi G, Donadio D, Parrinello M (2007) Canonical sampling through velocity rescaling. J Chem Phys 126: 014101

Cakmak FP, Choi S, Meyer MO, Bevilacqua PC, Keating CD (2020) Prebiotically-relevant low polyion multivalency can improve functionality of membraneless compartments. Nat Commun 11

Case DA, Aktulga HM, Belfon K, Ben-Shalom I, Berryman JT, Brozell SR, Cerutti DS, Cheatham TE, Cisneros GA, Cruzeiro VWD, 2022. Amber 2022. University of California, San Francisco.

Cheng T, Zhao Y, Li X, Lin F, Xu Y, Zhang X, Li Y, Wang R, Lai L (2007) Computation of octanol-water partition coefficients by guiding an additive model with knowledge. J Chem Inf Model 47: 2140–2148

Chong PA, Vernon RM, Forman-Kay JD (2018) RGG/RG Motif Regions in RNA Binding and Phase Separation. J Mol Biol 430: 4650–4665

Chong Y, Kleinhammes A, Tang P, Xu Y, Wu Y (2015) Dominant Alcohol-Protein Interaction via Hydration-Enabled Enthalpy-Driven Binding Mechanism. J Phys Chem B 119: 5367–5375

Dai B, Zhong T, Chen ZX, Chen W, Zhang N, Liu XL, Wang LQ, Chen J, Liang Y (2021) Myricetin slows liquid-liquid phase separation of Tau and activates ATG5-dependent autophagy to suppress Tau toxicity. J Biol Chem 297: 101222

Daigle JG, Lanson NA, Jr., Smith RB, Casci I, Maltare A, Monaghan J, Nichols CD, Kryndushkin D, Shewmaker F, Pandey UB (2013) RNA-binding ability of FUS regulates neurodegeneration, cytoplasmic mislocalization and incorporation into stress granules associated with FUS carrying ALS-linked mutations. Hum Mol Genet 22: 1193–1205

Das A, Mukhopadhyay C (2009) Urea-mediated protein denaturation: a consensus view. J Phys Chem B 113: 12816–12824

Deng H, Gao K, Jankovic J (2014) The role of FUS gene variants in neurodegenerative diseases. Nat Rev Neurol 10: 337–348

Düster R, Kaltheuner IH, Schmitz M, Geyer M (2021) 1,6-Hexanediol, commonly used to dissolve liquid–liquid phase separated condensates, directly impairs kinase and phosphatase activities. Journal of Biological Chemistry 296: 100260

Fuller GG, Han T, Freeberg MA, Moresco JJ, Ghanbari Niaki A, Roach NP, Yates JR, 3rd, Myong S, Kim JK (2020) RNA promotes phase separation of glycolysis enzymes into yeast G bodies in hypoxia. Elife 9

Gowers RJ, Linke M, Barnoud J, Reddy T, Melo MN, Seyler SL, Domanski JJ, Dotson DL, Buchoux S, Kenney IM et al, 2016. MDAnalysis: A Python Package for the Rapid Analysis of Molecular Dynamics Simulations, SciPy.

Gu J, Zhou X, Sutherland L, Kato M, Jaczynska K, Rizo J, McKnight SL (2023) Oxidative regulation of TDP-43 self-association by a β-to-α conformational switch. Proceedings of the National Academy of Sciences 120

Harris RK, Becker ED, Cabral De Menezes SM, Goodfellow R, Granger P (2001) NMR nomenclature. Nuclear spin properties and conventions for chemical shifts(IUPAC Recommendations 2001). Pure and Applied Chemistry 73: 1795–1818

He X, Man VH, Yang W, Lee TS, Wang J (2020) A fast and high-quality charge model for the next generation general AMBER force field. J Chem Phys 153: 114502

Hedtfeld M, Dammers A, Koerner C, Musacchio A (2024) A validation strategy to assess the role of phase separation as a determinant of macromolecular localization. Molecular Cell 84: 1783–1801.e1787

Hofweber M, Hutten S, Bourgeois B, Spreitzer E, Niedner-Boblenz A, Schifferer M, Ruepp MD, Simons M, Niessing D, Madl T et al (2018) Phase Separation of FUS Is Suppressed by Its Nuclear Import Receptor and Arginine Methylation. Cell 173: 706–719 e713

Hopkins CW, Le Grand S, Walker RC, Roitberg AE (2015) Long-Time-Step Molecular Dynamics through Hydrogen Mass Repartitioning. J Chem Theory Comput 11: 1864–1874

Itoh Y, Iida S, Tamura S, Nagashima R, Shiraki K, Goto T, Hibino K, Ide S, Maeshima K (2021) 1,6-hexanediol rapidly immobilizes and condenses chromatin in living human cells. Life Sci Alliance 4: e202001005

Jaggi RD, Franco-Obregon A, Muhlhausser P, Thomas F, Kutay U, Ensslin K (2003) Modulation of nuclear pore topology by transport modifiers. Biophys J 84: 665–670

Jiang S, Fagman JB, Chen C, Alberti S, Liu B (2020) Protein phase separation and its role in tumorigenesis. Elife 9: e60264

Kato M, McKnight SL (2018) A Solid-State Conceptualization of Information Transfer from Gene to Message to Protein. Annu Rev Biochem 87: 351–390

Klein IA, Boija A, Afeyan LK, Hawken SW, Fan M, Dall’Agnese A, Oksuz O, Henninger JE, Shrinivas K, Sabari BR et al (2020) Partitioning of cancer therapeutics in nuclear condensates. Science 368: 1386–1392

Kroschwald S, Maharana S, Mateju D, Malinovska L, Nuske E, Poser I, Richter D, Alberti S (2015) Promiscuous interactions and protein disaggregases determine the material state of stress-inducible RNP granules. Elife 4: e06807

Kroschwald S, Maharana S, Simon A (2017) Hexanediol: a chemical probe to investigate the material properties of membrane-less compartments. Matters

Levone BR, Lenzken SC, Antonaci M, Maiser A, Rapp A, Conte F, Reber S, Mechtersheimer J, Ronchi AE, Muhlemann O et al (2021) FUS-dependent liquid-liquid phase separation is important for DNA repair initiation. J Cell Biol 220: e202008030

Li S, Yoshizawa T, Yamazaki R, Fujiwara A, Kameda T, Kitahara R (2021) Pressure and Temperature Phase Diagram for Liquid-Liquid Phase Separation of the RNA-Binding Protein Fused in Sarcoma. J Phys Chem B 125: 6821–6829

Lin Y, Mori E, Kato M, Xiang S, Wu L, Kwon I, McKnight SL (2016) Toxic PR Poly-Dipeptides Encoded by the C9orf72 Repeat Expansion Target LC Domain Polymers. Cell 167: 789–802 e712

Liu X, Jiang S, Ma L, Qu J, Zhao L, Zhu X, Ding J (2021) Time-dependent effect of 1,6-hexanediol on biomolecular condensates and 3D chromatin organization. Genome Biol 22: 230

Loughlin FE, Lukavsky PJ, Kazeeva T, Reber S, Hock EM, Colombo M, Von Schroetter C, Pauli P, Clery A, Muhlemann O et al (2019) The Solution Structure of FUS Bound to RNA Reveals a Bipartite Mode of RNA Recognition with Both Sequence and Shape Specificity. Mol Cell 73: 490–504 e496

Lu J, Qian J, Xu Z, Yin S, Zhou L, Zheng S, Zhang W (2021) Emerging Roles of Liquid-Liquid Phase Separation in Cancer: From Protein Aggregation to Immune-Associated Signaling. Front Cell Dev Biol 9: 631486

Luo Y, Roux B (2009) Simulation of Osmotic Pressure in Concentrated Aqueous Salt Solutions. The Journal of Physical Chemistry Letters 1: 183–189

Lyon AS, Peeples WB, Rosen MK (2021) A framework for understanding the functions of biomolecular condensates across scales. Nat Rev Mol Cell Biol 22: 215–235

Martin EW, Holehouse AS, Peran I, Farag M, Incicco JJ, Bremer A, Grace CR, Soranno A, Pappu RV, Mittag T (2020) Valence and patterning of aromatic residues determine the phase behavior of prion-like domains. Science 367: 694–699

Martin EW, Mittag T (2018) Relationship of Sequence and Phase Separation in Protein Low-Complexity Regions. Biochemistry 57: 2478–2487

Mitrea DM, Mittasch M, Gomes BF, Klein IA, Murcko MA (2022) Modulating biomolecular condensates: a novel approach to drug discovery. Nat Rev Drug Discov 21: 841–862

Miyawaki O, Tatsuno M (2011) Thermodynamic analysis of alcohol effect on thermal stability of proteins. J Biosci Bioeng 111: 198–203

Monahan Z, Ryan VH, Janke AM, Burke KA, Rhoads SN, Zerze GH, O’Meally R, Dignon GL, Conicella AE, Zheng W et al (2017) Phosphorylation of the FUS low-complexity domain disrupts phase separation, aggregation, and toxicity. EMBO J 36: 2951–2967

Murthy AC, Dignon GL, Kan Y, Zerze GH, Parekh SH, Mittal J, Fawzi NL (2019) Molecular interactions underlying liquid-liquid phase separation of the FUS low-complexity domain. Nat Struct Mol Biol 26: 637–648

Murthy AC, Tang WS, Jovic N, Janke AM, Seo DH, Perdikari TM, Mittal J, Fawzi NL (2021) Molecular interactions contributing to FUS SYGQ LC-RGG phase separation and co-partitioning with RNA polymerase II heptads. Nat Struct Mol Biol 28: 923–935

Nakata N, Okamoto R, Sumi T, Koga K, Morita T, Imamura H (2023) Molecular mechanism of the common and opposing cosolvent effects of fluorinated alcohol and urea on a coiled coil protein. Protein Sci 32: e4763

Naumann M, Pal A, Goswami A, Lojewski X, Japtok J, Vehlow A, Naujock M, Gunther R, Jin M, Stanslowsky N et al (2018) Impaired DNA damage response signaling by FUS-NLS mutations leads to neurodegeneration and FUS aggregate formation. Nat Commun 9: 335

Owen I, Yee D, Wyne H, Perdikari TM, Johnson V, Smyth J, Kortum R, Fawzi NL, Shewmaker F (2021) The oncogenic transcription factor FUS-CHOP can undergo nuclear liquid-liquid phase separation. J Cell Sci 134: jcs258578

Ozdilek BA, Thompson VF, Ahmed NS, White CI, Batey RT, Schwartz JC (2017) Intrinsically disordered RGG/RG domains mediate degenerate specificity in RNA binding. Nucleic Acids Res 45: 7984–7996

Ozguney B, Mohanty P, Mittal J (2024) RNA binding tunes the conformational plasticity and intradomain stability of TDP-43 tandem RNA recognition motifs. Biophysical Journal 123: 3844–3855

Palmer AG, Kroenke CD, Patrick Loria J (2001) Nuclear Magnetic Resonance Methods for Quantifying Microsecond-to-Millisecond Motions in Biological Macromolecules. In: Methods in Enzymology, pp. 204–238. Elsevier:

Parrinello M, Rahman A (1980) Crystal Structure and Pair Potentials: A Molecular-Dynamics Study. Physical Review Letters 45: 1196–1199

Pastor RW (1994) Techniques and Applications of Langevin Dynamics Simulations. In: The Molecular Dynamics of Liquid Crystals, Luckhurst G.R., Veracini C.A. (eds.) pp. 85–138. Springer Netherlands: Dordrecht

Patel A, Lee HO, Jawerth L, Maharana S, Jahnel M, Hein MY, Stoynov S, Mahamid J, Saha S, Franzmann TM et al (2015) A Liquid-to-Solid Phase Transition of the ALS Protein FUS Accelerated by Disease Mutation. Cell 162: 1066–1077

Perry SL, Leon L, Hoffmann KQ, Kade MJ, Priftis D, Black KA, Wong D, Klein RA, Pierce CF, Margossian KO et al (2015) Chirality-selected phase behaviour in ionic polypeptide complexes. Nat Commun 6

Rhine K, Dasovich M, Yoniles J, Badiee M, Skanchy S, Ganser LR, Ge Y, Fare CM, Shorter J, Leung AKL et al (2022) Poly(ADP-ribose) drives condensation of FUS via a transient interaction. Mol Cell 82: 969–985 e911

Ribbeck K, Gorlich D (2002) The permeability barrier of nuclear pore complexes appears to operate via hydrophobic exclusion. EMBO J 21: 2664–2671

Roden C, Gladfelter AS (2021) RNA contributions to the form and function of biomolecular condensates. Nat Rev Mol Cell Biol 22: 183–195

Roe DR, Cheatham TE, 3rd (2013) PTRAJ and CPPTRAJ: Software for Processing and Analysis of Molecular Dynamics Trajectory Data. J Chem Theory Comput 9: 3084–3095

Rog O, Kohler S, Dernburg AF (2017) The synaptonemal complex has liquid crystalline properties and spatially regulates meiotic recombination factors. Elife 6: e21455

Romero CM, Páez MS, Arteaga JC, Romero MA, Negrete F (2007) Effect of temperature on the volumetric properties of dilute aqueous solutions of 1,2-hexanediol, 1,5-hexanediol, 1,6-hexanediol, and 2,5-hexanediol. The Journal of Chemical Thermodynamics 39: 1101–1109

Ryan VH, Fawzi NL (2019) Physiological, Pathological, and Targetable Membraneless Organelles in Neurons. Trends Neurosci 42: 693–708

Sabari BR, Dall’Agnese A, Boija A, Klein IA, Coffey EL, Shrinivas K, Abraham BJ, Hannett NM, Zamudio AV, Manteiga JC et al (2018) Coactivator condensation at super-enhancers links phase separation and gene control. Science 361

Sama RR, Ward CL, Bosco DA (2014) Functions of FUS/TLS from DNA repair to stress response: implications for ALS. ASN Neuro 6

Sato K, Sakai M, Ishii A, Maehata K, Takada Y, Yasuda K, Kotani T (2022) Identification of embryonic RNA granules that act as sites of mRNA translation after changing their physical properties. iScience 25: 104344

Schmidt HB, Jaafar ZA, Wulff BE, Rodencal JJ, Hong K, Aziz-Zanjani MO, Jackson PK, Leonetti MD, Dixon SJ, Rohatgi R et al (2022) Oxaliplatin disrupts nucleolar function through biophysical disintegration. Cell Reports 41: 111629

Shi M, You K, Chen T, Hou C, Liang Z, Liu M, Wang J, Wei T, Qin J, Chen Y et al (2021) Quantifying the phase separation property of chromatin-associated proteins under physiological conditions using an anti-1,6-hexanediol index. Genome Biol 22: 229

Shin Y, Brangwynne CP (2017) Liquid phase condensation in cell physiology and disease. Science 357

Shirts MR, Klein C, Swails JM, Yin J, Gilson MK, Mobley DL, Case DA, Zhong ED (2017) Lessons learned from comparing molecular dynamics engines on the SAMPL5 dataset. J Comput Aided Mol Des 31: 147–161

Shulga N, Goldfarb DS (2003) Binding dynamics of structural nucleoporins govern nuclear pore complex permeability and may mediate channel gating. Mol Cell Biol 23: 534–542

Snead WT, Gladfelter AS (2019) The Control Centers of Biomolecular Phase Separation: How Membrane Surfaces, PTMs, and Active Processes Regulate Condensation. Mol Cell 76: 295–305

Sousa da Silva AW, Vranken WF (2012) ACPYPE - AnteChamber PYthon Parser interfacE. BMC Res Notes 5: 367

Sutton EC, DeRose VJ (2021) Early nucleolar responses differentiate mechanisms of cell death induced by oxaliplatin and cisplatin. J Biol Chem 296: 100633

Thandapani P, O’Connor TR, Bailey TL, Richard S (2013) Defining the RGG/RG motif. Mol Cell 50: 613–623

Trnka F, Hoffmann C, Wang H, Sansevrino R, Rankovic B, Rost BR, Schmitz D, Schmidt HB, Milovanovic D (2021) Aberrant Phase Separation of FUS Leads to Lysosome Sequestering and Acidification. Front Cell Dev Biol 9

Tulpule A, Guan J, Neel DS, Allegakoen HR, Lin YP, Brown D, Chou YT, Heslin A, Chatterjee N, Perati S et al (2021) Kinase-mediated RAS signaling via membraneless cytoplasmic protein granules. Cell 184: 2649–2664 e2618

Uechi H, Sridharan S, Nijssen J, Bilstein J, Iglesias-Artola JM, Kishigami S, Casablancas-Antras V, Poser I, Martinez EJ, Boczek E et al, 2019. Small molecule modulation of a redox-sensitive stress granule protein dissolves stress granules with beneficial outcomes for familial amyotrophic lateral sclerosis models. Cold Spring Harbor Laboratory.

Ulianov SV, Velichko AK, Magnitov MD, Luzhin AV, Golov AK, Ovsyannikova N, Kireev, II, Gavrikov AS, Mishin AS, Garaev AK et al (2021) Suppression of liquid-liquid phase separation by 1,6-hexanediol partially compromises the 3D genome organization in living cells. Nucleic Acids Res 49: 10524–10541

Wake N, Weng SL, Zheng T, Wang SH, Kirilenko V, Mittal J, Fawzi NL (2024) Expanding the molecular grammar of polar residues and arginine in FUS prion-like domain phase separation and aggregation. bioRxiv: doi: 10.1101/2024.1102.1115.580391

Wang J, Choi JM, Holehouse AS, Lee HO, Zhang X, Jahnel M, Maharana S, Lemaitre R, Pozniakovsky A, Drechsel D et al (2018) A Molecular Grammar Governing the Driving Forces for Phase Separation of Prion-like RNA Binding Proteins. Cell 174: 688–699 e616

Wheeler JR, Matheny T, Jain S, Abrisch R, Parker R (2016) Distinct stages in stress granule assembly and disassembly. Elife 5

Wolozin B, Ivanov P (2019) Stress granules and neurodegeneration. Nat Rev Neurosci 20: 649–666

Yang P, Mathieu C, Kolaitis R-M, Zhang P, Messing J, Yurtsever U, Yang Z, Wu J, Li Y, Pan Q et al (2020) G3BP1 Is a Tunable Switch that Triggers Phase Separation to Assemble Stress Granules. Cell 181: 325–345.e328

Zheng W, Dignon GL, Jovic N, Xu X, Regy RM, Fawzi NL, Kim YC, Best RB, Mittal J (2020) Molecular Details of Protein Condensates Probed by Microsecond Long Atomistic Simulations. J Phys Chem B 124: 11671–11679

