## Supplemental Figurs and Table for "Molecular insights into the effect of hexanediol on FUS phase separation"

†These authors contributed equally

**Table of Contents**

| | Page |
|---|---|
| Appendix Figure S1 | 2 |
| Appendix Figure S2 | 3 |
| Appendix Figure S3 | 4 |
| Appendix Figure S4 | 5 |
| Appendix Figure S5 | 6 |
| Appendix Figure S6 | 7 |
| Appendix Figure S7 | 8 |
| Appendix Figure S8 | 9 |
| Appendix Figure S9 | 10 |
| Appendix Figure S10 | 11 |
| Appendix Figure S11 | 12 |
| Appendix Table S1 | 13 |
| Appendix Table S2 | 13 |

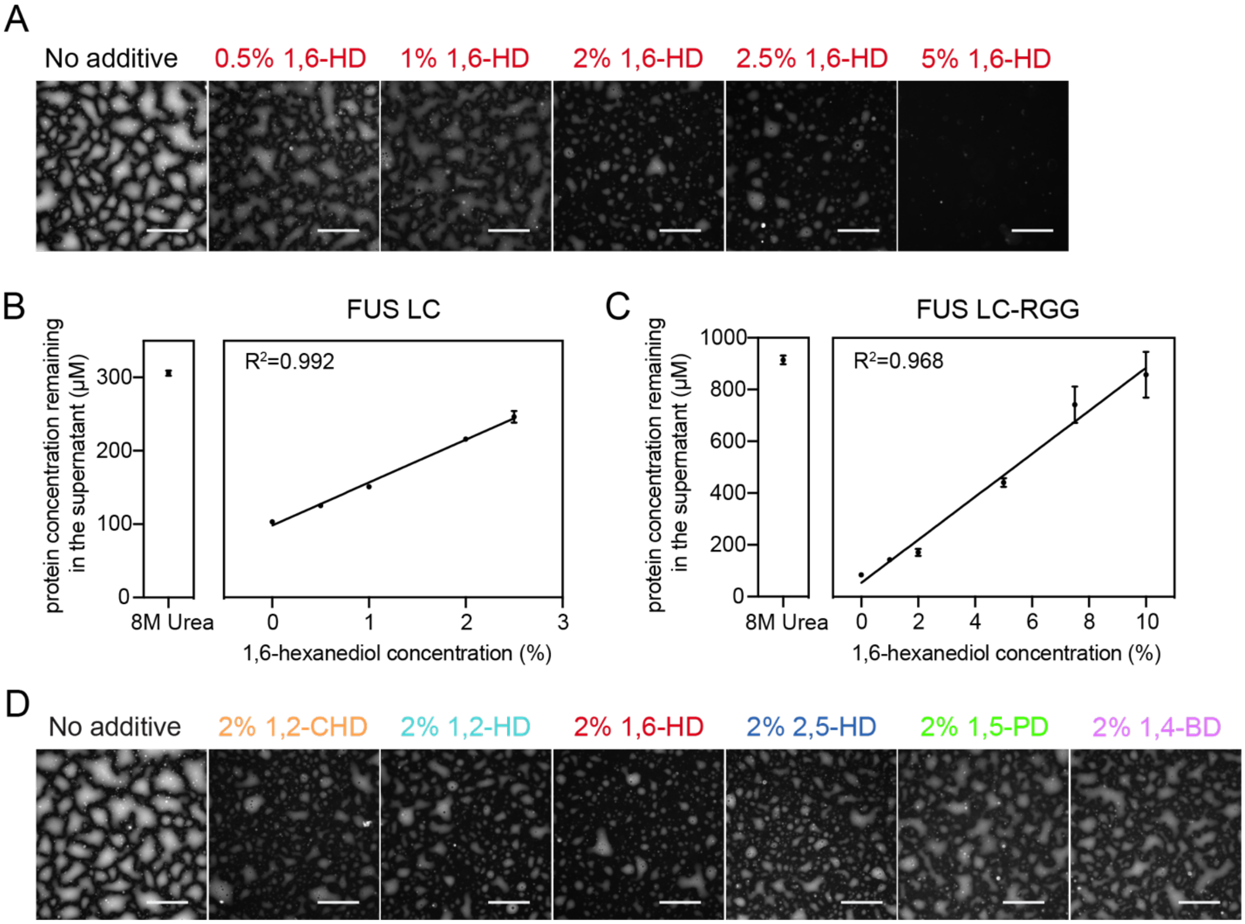


**Appendix Figure S1. Alkanediols disrupt FUS LC and LC-RGG1 phase separation with a linear dependence on concentration up to 10%.**

A) Fluorescence microscopy images of 300 μM FUS LC after treatment with 0-5% w/v 1,6-Hexanediol and 20 μM ThT, which serves as a (weakly) fluorescent dye that partitions to or marks condensates. Scale bars represent 100 μm.

B) Saturation concentration of FUS LC in the presence of 0-2.5% 1,6-HD.

C) Saturation concentration of FUS LC-RGG in the presence of 0-10% 1,6-HD.

D) Fluorescence microscopy images of 300 μM FUS LC after treatment with 5% w/v aliphatic alcohols and 20 μM ThT. Scale bars represent 100 μm.

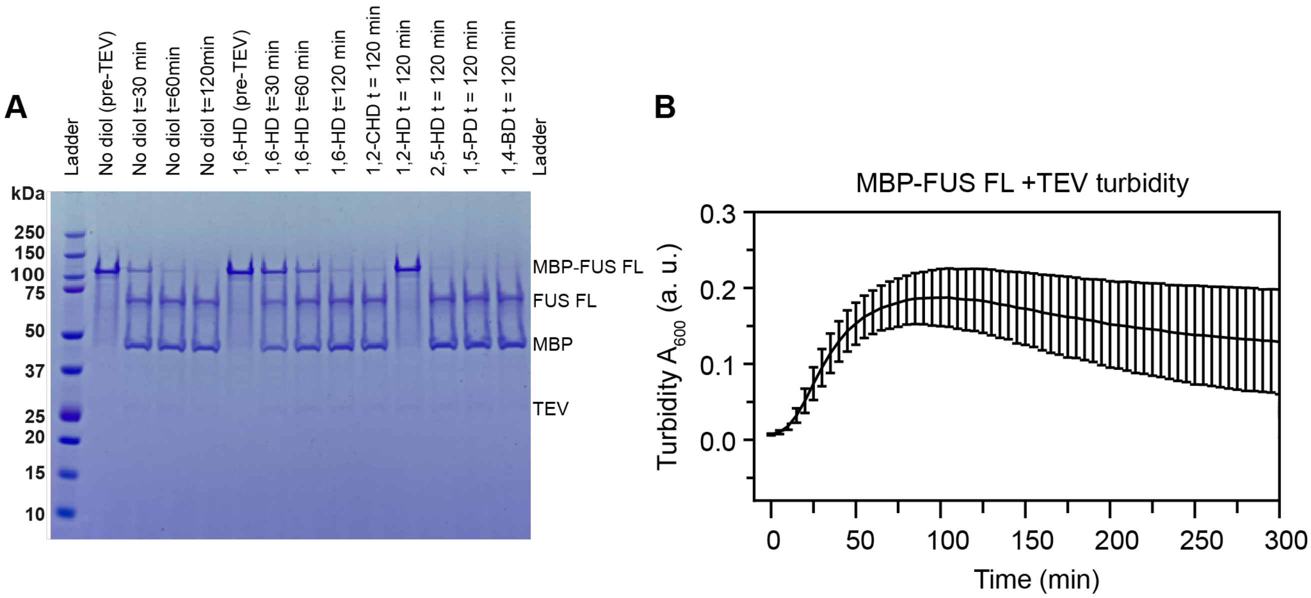


**Appendix Figure S2. Impact of alkanediols on TEV cleavage of MBP tag from MBP-FUS FL**

A) Analysis of TEV-mediated cleavage of FUS full-length in the presence of different alkanediols by SDS-PAGE. These data show that cleaving is slightly slowed by 1,6-HD (center) and is completely inhibited (at 120 minutes) by 1,2-HD. Therefore, all experiments in the main text with FUS full-length were performed where the cleaving is performed first in buffer without alkanediols, followed by addition of alkanediols (and RNA, if applicable).

B) TEV-mediated cleavage was monitored using a turbidity assay. FUS FL phase separates readily when the MBP tag is cleaved off.

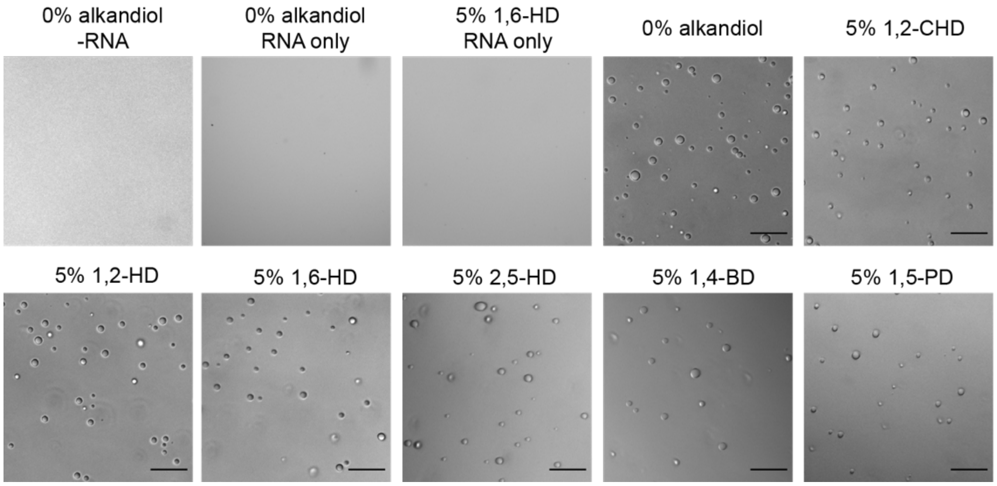


**Appendix Figure S3. Alkanediols do not significantly affect FUS RGG3-RNA condensates.**
Droplets were made in 50 mM MES pH 5.5, 150 mM NaCl, 5% co-solvent, 100 μM FUS RGG3, 0.142 mg/ml polyadenylic acid (polyA) RNA and were imaged after 1 hour of incubation in room temperature. Scale bars represent 50 μm.

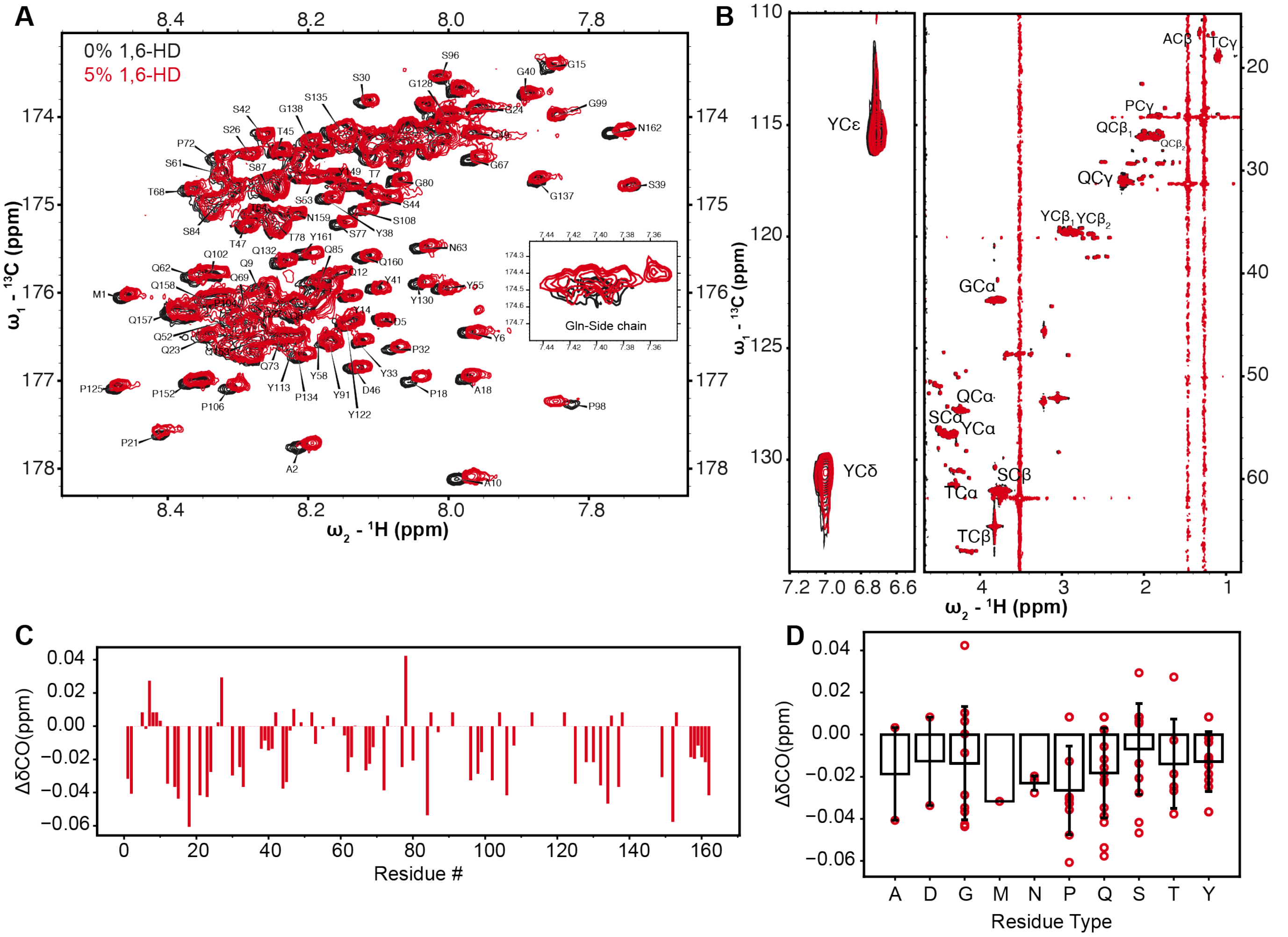


**Appendix Figure S4. 1,6-hexanediol alters FUS chemical environment.**

A) $^{1}H$-$^{13}C$ two-dimensional spectral overlay with (red) and without (black) 5% w/v 1,6-hexanediol derived from a triple resonance HNCO experiment for 100 μM $^{15}N$, $^{13}C$ labelled FUS LC, 50 mM MES pH 5.5, 150mM NaCl. Inset corresponds to the $^{13}C$ carbonyl position in the side-chain of glutamine (note, the chemical shift position is aliased).

B) $^{13}C$ HSQC of 100 μM $^{15}N$, $^{13}C$ labelled FUS LC, 50 mM MES pH 5.5, 150mM NaCl with (red) and without (black) 1,6-hexanediol.

C) 1,6-hexanediol induced chemical shift deviations of $^{13}CO$ at each assigned backbone position.

D) $^{13}CO$ chemical shift deviations binned by residue type. Individual residues are plotted as red marks. Bar plots represent mean and standard deviation among all chemical shifts of each residue type.

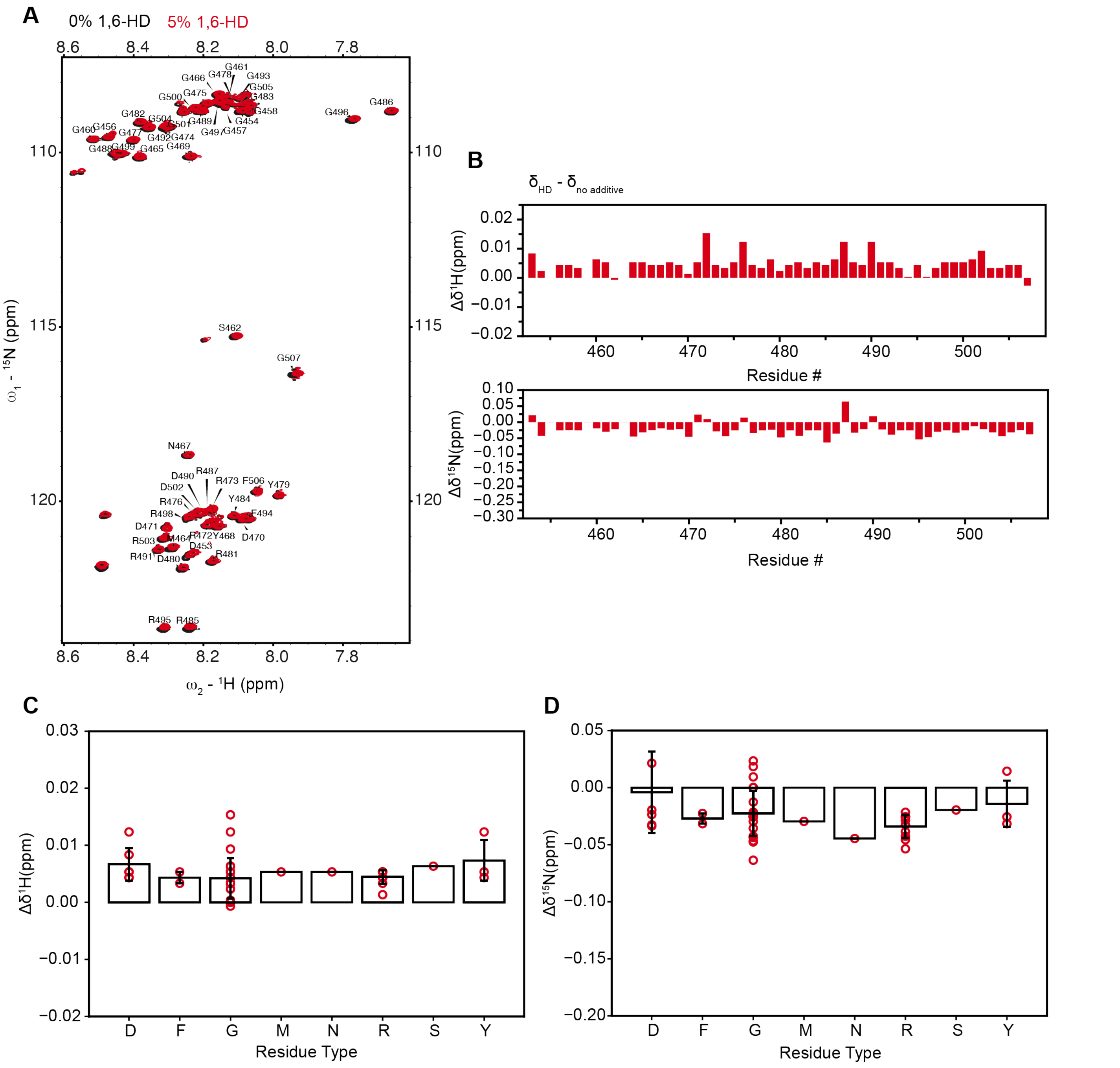


**Appendix Figure S5. Effect of 1,6-hexanediol on FUS RGG3 NMR spectra**

A) $^{1}$H-$^{15}$N heteronuclear single quantum coherence (HSQC) spectrum of 100 µM FUS RGG3 with (red) and without (black) 5% 1,6-hexanediol.

B) $^{1}$H (top) and $^{15}$N chemical shift perturbations caused by 5% 1,6-hexanediol of FUS RGG3.

C-D) Chemical shifts binned by residue type. Individual chemical shifts are plotted as blue marks. Bar plots represent mean and s.d. among all chemical shifts of each residue type.

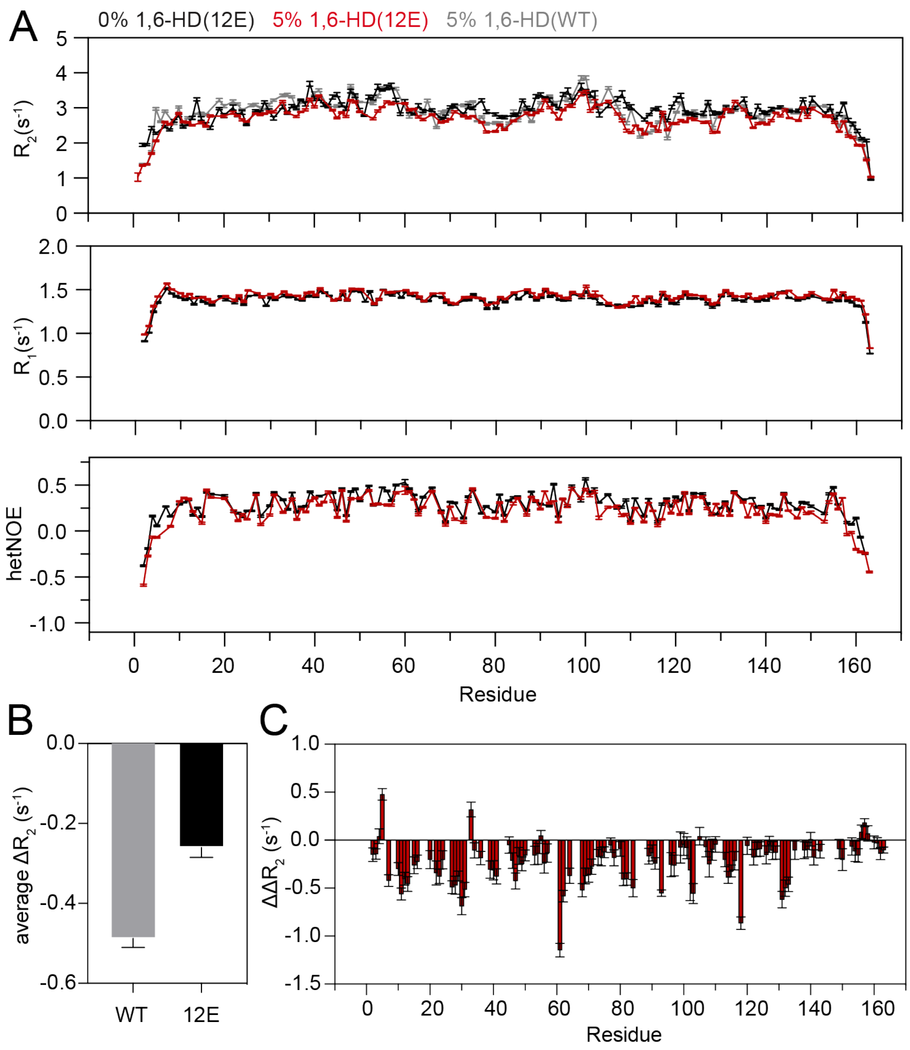


**Appendix Figure S6. Less change due to 1,6-hexanediol in the molecular motions of the phosphomimetic mutant FUS LC 12E that does not undergo phase separation compared to the wild-type.**

A) NMR spin relaxation parameters $^{15}N$ $R_2$, $^{15}N$ $R_1$ and ($^1H$) $^{15}N$ heteronuclear NOE values measured at 850 MHz $^1H$ frequency for the phase-separation resistant FUS SYGQ LC 12E phosphomimetic variant with twelve serine-to-glutamate substitutions across the entire domain.

B) Average $\Delta R_2$ values for FUS LC WT or 12E variant resulting from the addition of 5% 1,6-HD. The addition of 5% 1,6-HD caused an approximately two-fold greater reduction in $\Delta R_2$ values for WT protein compared to 12E variant.

C) Residue-specific $\Delta\Delta R_2$ values show reductions in $\Delta R_2$ across FUS LC residues when comparing WT to 12E variant. Error bars represent standard deviations. These observations suggest that 1,6-HD induced reduction in wild-type $R_2$ are partially due to disruption of contacts that are already disrupted in this phosphomimetic variant.

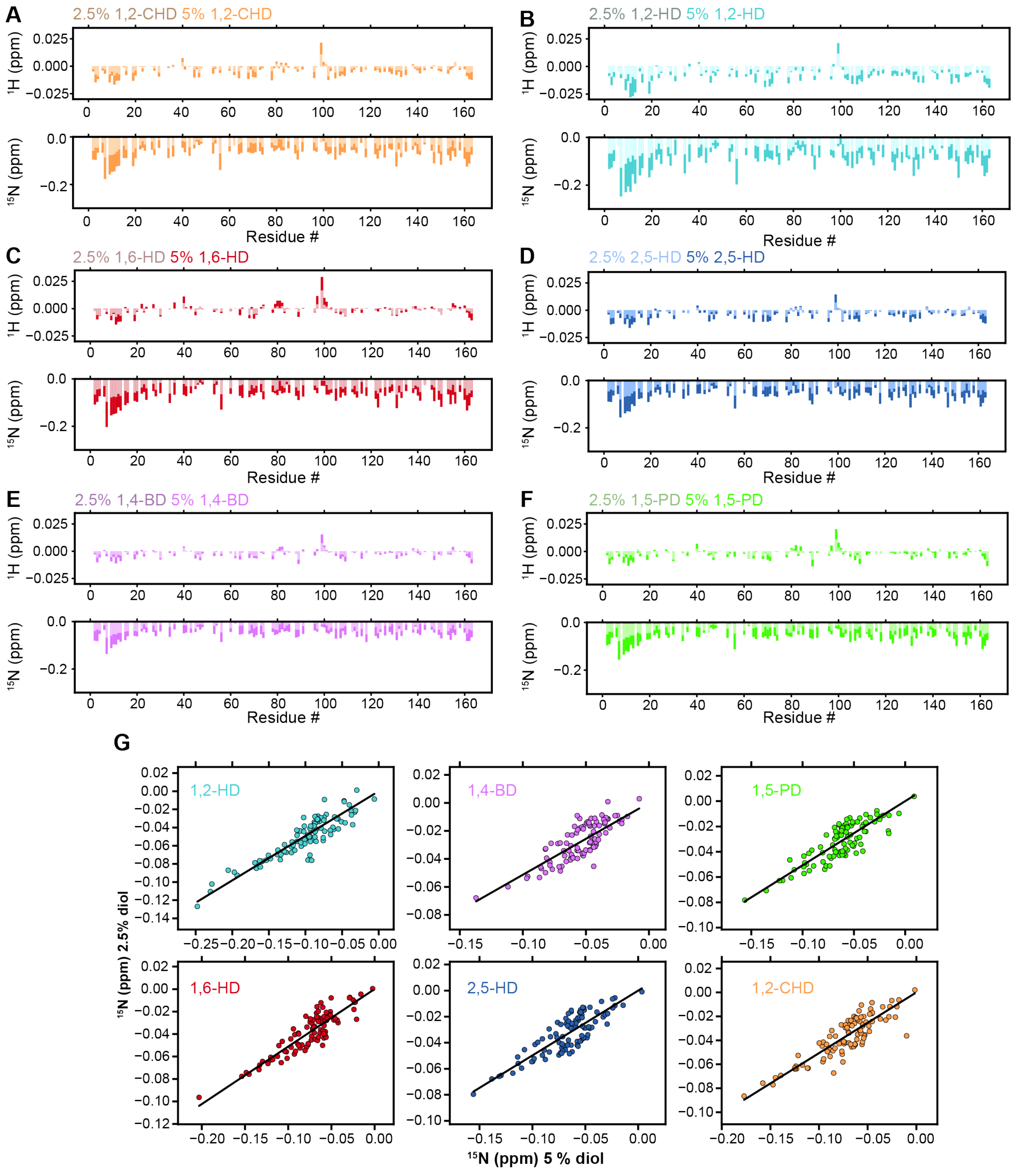


**Appendix Figure S7. Alkandediols quantitatively affect the chemical environment of FUS LC.**

A-F) Chemical shift deviations for FUS LC in the presence of different diols at 2.5% or 5% w/v in 50 mM MES pH 5.5, 150 mM NaCl.

G) Correlation of $^{15}N$ chemical shift deviations for 2.5% and 5% diol.

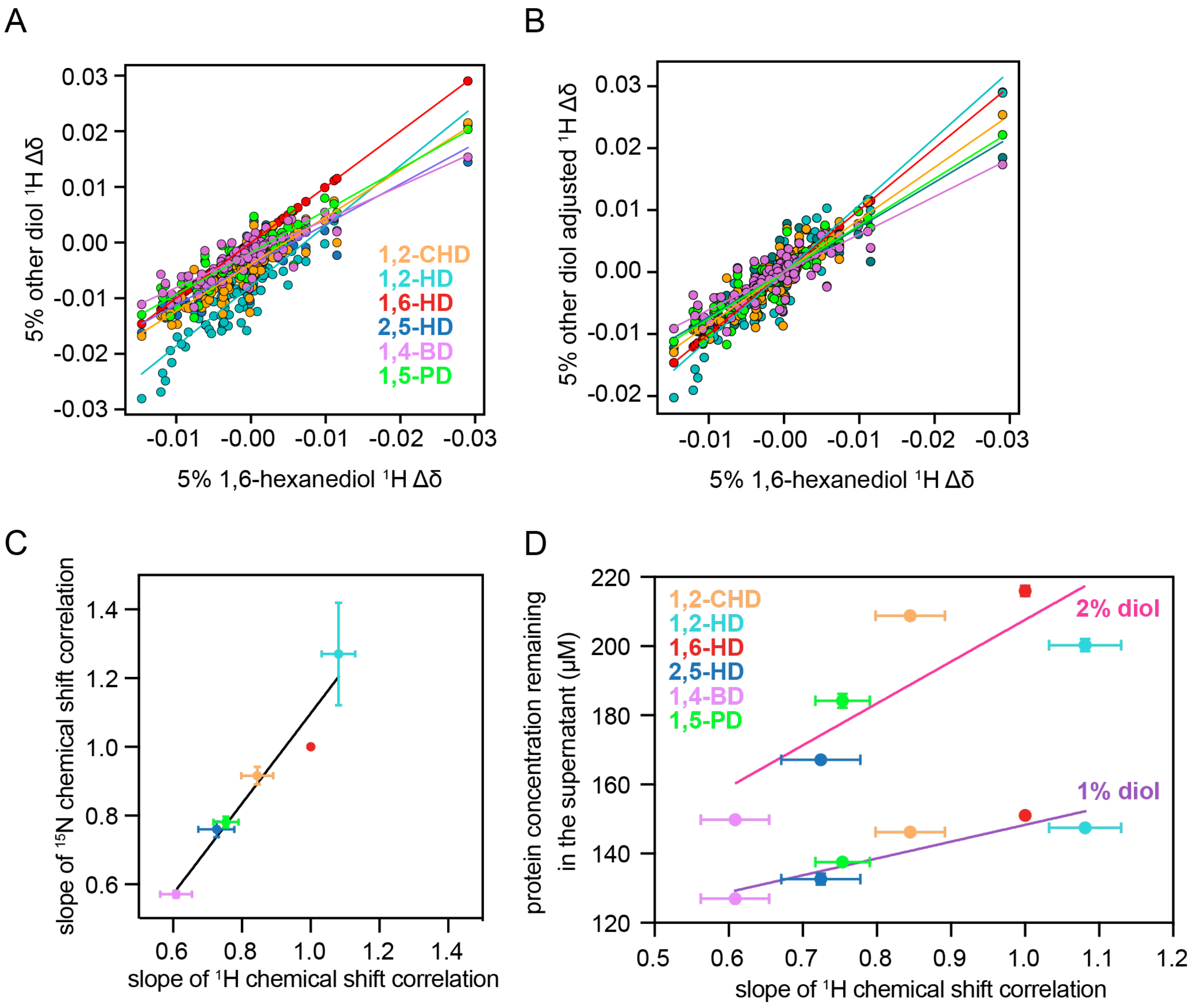


**Appendix Figure S8. FUS LC $^{1}$H chemical shift perturbations report on the changes in chemical environment due to alkanediols similarly to the $^{15}$N chemical shift perturbations.**

A) Comparison of $^{1}$H Δδ in 5% 1,6-hexanediol with every other co-solvent. Fit comparing 1,6-HD to other alkanediols do not pass through the origin because $^{1}$H chemical shift referencing is minimally but noticeably affected by the change in solvation environment of DSS by the alkanediol addition (in other words, DSS reference compound experiences differential chemical shift perturbations in the presence of each alkanediol), which is not surprising for the high gyromagnetic ratio $^{1}$H nucleus upon addition of 10% w/w amounts of co-solvent. However, these small constant offsets do not affect the residue-specific chemical shift perturbations also observed in $^{1}$H CSPs (see B, below).

B) $^{1}$H Δδ shifted by a constant offset for each alkanediol so that all the fittings are y=mx and pass through (0,0).

C) Correlation between slope of the $^{1}$H and $^{15}$N chemical shift perturbations normalized to 1,6-HD demonstrating that both $^{1}$H and $^{15}$N chemical shift perturbations report on the same change in chemical environment.

D) Slope extracted from the correlation presented above versus protein remaining in the supernatant shows that the $^{1}$H chemical shift differences induced by each diol are also correlated with the capacity of FUS LC to phase separate in each condition (PCC = 0.92 at 1% diol, PCC = 0.84 at 2% diol). Solid lines represent linear fits.

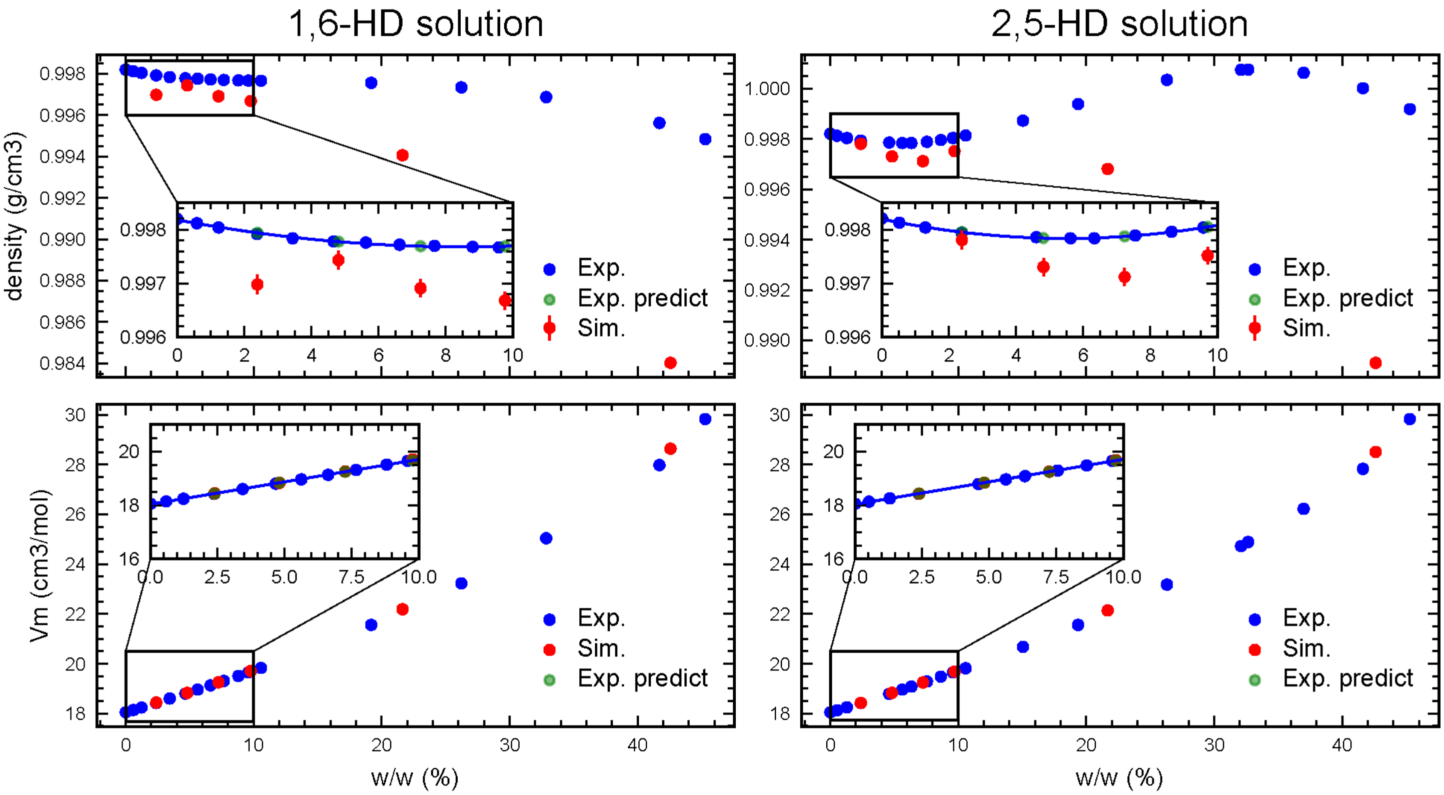


**Appendix Figure S9. Simulated values of hexanediol solutions compared to experimental data for parameter validation.**

The experimental predicted values were extrapolated from the reference (Romero *et al*, 2007).

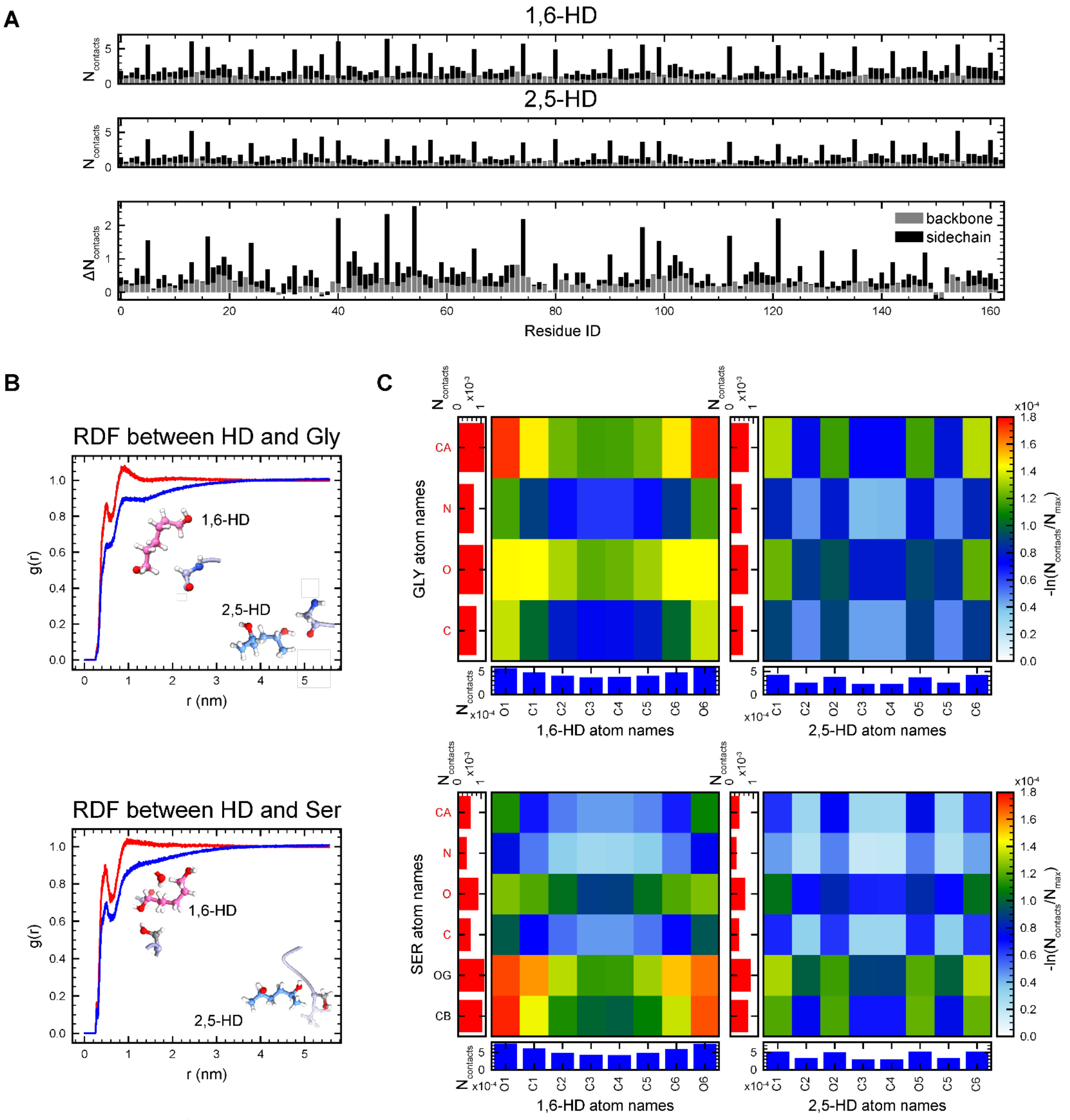


**Appendix Figure S10. 1,6- and 2,5-hexanediol directly interact with protein.**

A) Protein-hexanediol contact profiles calculated from atomistic simulations of single chain FUS LC with 5% 1,6-HD (top), 5% 2,5-HD (middle), and the difference (1,6-HD minus 2,5-HD, bottom).

B) Radial distribution functions (RDF, g(r)) between the center of mass (COM) of backbone heavy atoms of Gly / sidechain heavy atoms of Ser and heavy atoms of hexanediols. The corresponding interaction snapshots are inset.

C) Atomistic contact maps of Gly / Ser with hexanediols, calculated from the atomistic simulations. The left column and the bottom row of each panel show the one-dimensional summation of corresponding atomistic contacts.

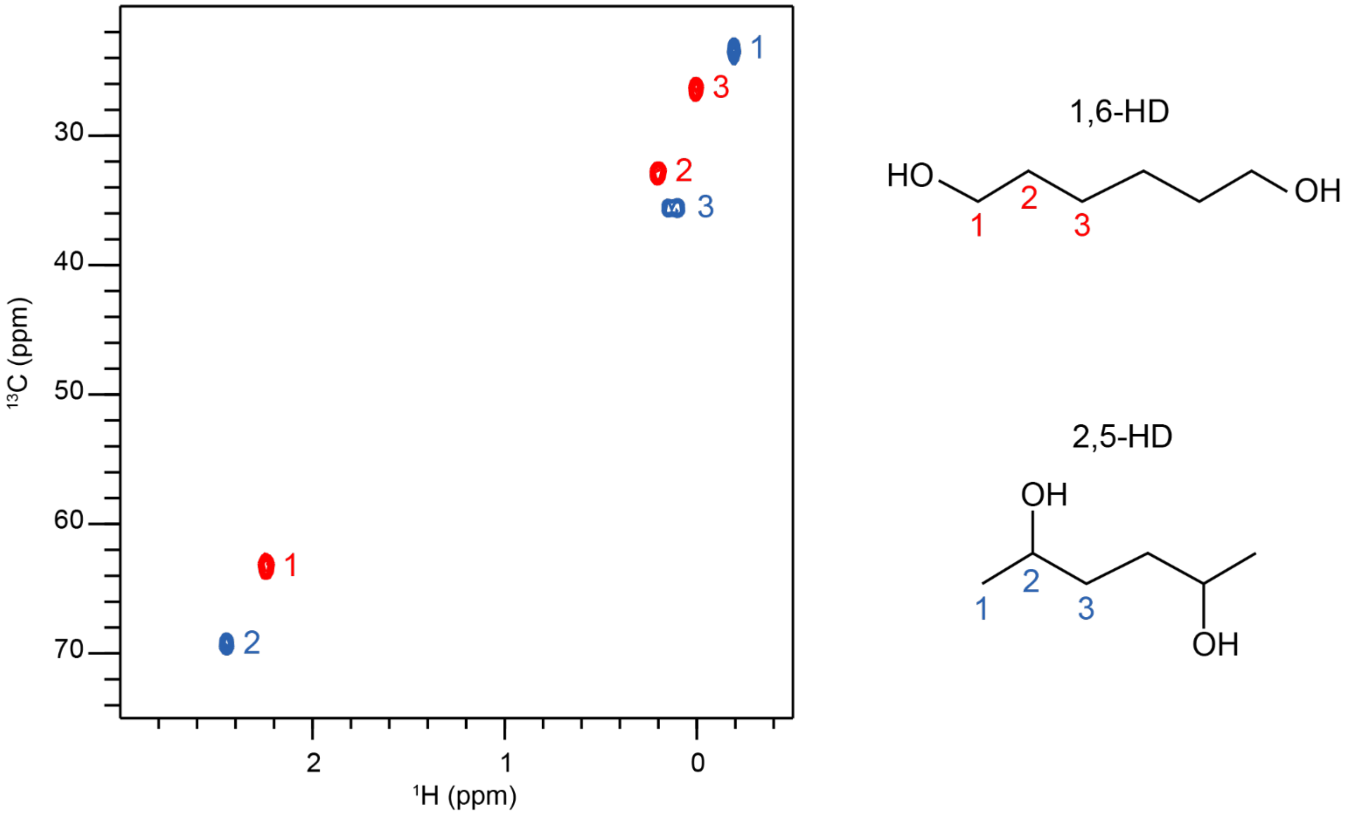


**Appendix Figure S11. $^{1}H$-$^{13}C$ HSQC spectra overlay of 1,6-HD and 2,5-HD.**
Two dimensional spectra of 5% hexanediol (at natural isotopic abundance of $^{13}C$) in 50 mM MES pH 7 provides information on the chemical environment of the carbon/hydrogen positions in hexanediol and shows how the bonding differences in the isomers affects the chemical environment.

**Appendix Table S1: Simulated hexanediol solutions densities for parameter validation.** Since data with identical simulated concentrations was not available, the experimental densities were extrapolated from reference (Romero *et al*, 2007).

| ρ (g/ml) | 2.4% | 4.8% | 7.2% | 9.7% | 21.7% | 42.6% |
|---|---|---|---|---|---|---|
| 1,6-HD | 0.996981 | 0.997434 | 0.996910 | 0.996681 | 0.994061 | 0.984024 |
| Exp | 0.997941 | 0.997777 | 0.997692 | 0.997690 | n. a. | n. a. |
| Error % | -0.0962% | -0.0343% | -0.0784% | -0.1011% | | |
| 2,5-HD | 0.997809 | 0.997307 | 0.99712 | 0.997518 | 0.996812 | 0.989105 |
| Exp | 0.997953 | 0.997843 | 0.997874 | 0.998050 | n. a. | n. a. |
| Error % | -0.0144% | -0.0537% | -0.0756% | -0.0533% | | |

**Appendix Table S2: Simulated hexanediol solutions molar volumes for parameter validation.** Since data with identical simulated concentrations was not available, the experimental molar volumes were extrapolated from reference (Romero *et al*, 2007).

| ρ (g/ml) | 2.4% | 4.8% | 7.2% | 9.7% | 21.7% | 42.6% |
|---|---|---|---|---|---|---|
| 1,6-HD | 19.254991 | 18.828569 | 19.254991 | 19.707106 | 22.189575 | 28.639371 |
| Exp | 18.431929 | 18.835278 | 19.2438 | 19.663544 | n. a. | n. a. |
| Error % | 0.0612% | -0.0356% | 0.0582% | 0.222% | | |
| 2,5-HD | 18.427631 | 18.832728 | 19.247844 | 19.67962 | 22.139689 | 28.50896 |
| Exp | 18.432009 | 18.83918 | 19.244728 | 19.659182 | n. a. | n. a. |
| Error % | -0.0238% | -0.0342% | 0.0162% | 0.104% | | |

Romero, CM.; Páez, MS; Arteaga, JC; Romero, MA; Negrete F, (2007) Effect of Temperature on the Volumetric Properties of Dilute Aqueous Solutions of 1,2-Hexanediol, 1,5-Hexanediol, 1,6-Hexanediol, and 2,5-Hexanediol. *J Chem Thermodyn 39* (8): 1101–1109.
